# From head to tail - Atomistic mechanism of long-range coupling from the cytosolic sensor domain to the selectivity filter in TREK K_2P_ channels

**DOI:** 10.1101/2023.10.06.561191

**Authors:** Berke Türkaydin, Marcus Schewe, Elena Riel, Friederike Schulz, Johann Biedermann, Thomas Baukrowitz, Han Sun

**Author notes:** These authors contributed equally to this work (Berke Türkaydin, Marcus Schewe).

## Abstract

The two-pore domain potassium (K_2P_) channels TREK-1 and TREK-2 link neuronal excitability to a variety of stimuli including mechanical force, lipids, temperature and phosphorylation. This regulation involves the C-terminus as a polymodal stimulus sensor and the selectivity filter (SF) as channel gate. Using crystallographic up- and down-state structures of TREK-2 as a template for full atomistic molecular dynamics simulations, we reveal that the SF in down-state undergoes inactivation via conformational changes at the S1 ion coordination site, while the up-state structure maintains a stable and conductive SF. This provides an atomistic understanding of the low channel activity previously assigned to the down state, but not evident from the crystal structure. Furthermore, by using (de-)phosphorylation mimics and chemically attaching lipid tethers to the proximal C-terminus (pCt), we confirm the hypothesis that moving the pCt towards the membrane induces the up-state. We also uncover two gating pathways by which movement of the pCt controls the stability (i.e. conductivity) of the filter gate. Together, these findings provide atomistic insights into the SF gating mechanism and the physiological regulation of TREK channels by phosphorylation.

## Introduction

TREK channels belong to the two-pore domain potassium (K_2P_) channels, which are responsible for generating background currents that help maintain the resting membrane potential below the threshold of depolarization^1–3^. They are widespread throughout the cardiovascular, central, and peripheral nervous systems^4–7^, playing key roles in diverse physiological processes and diseases such as anesthesia, ischemia, epilepsy and depression. Loss-of-function mutations in these channels can lead to severe pathologies^8–11^. Structurally, TREK channels can exist as homo- or heterodimeric forms, each subunit comprising four membrane-spanning domains (M1 to M4), an extracellular loop domain (EC1/EC2), two pore-forming domains (P1 and P2) that built up a pseudo-tetrameric selectivity filter (SF) (Fig. 1a), and a long cytoplasmic C-terminus (Ct) connected to the distal M4 helix. While no full-length atomic structure of any K_2P_ channel has been resolved, X-ray structures of TREK-1, TREK-2 and TRAAK have been determined in different N- and C-terminal truncated forms^12–17^. These structures imply flexibility in the long Ct, although AlphaFold2^18^ suggested a helical conformation for most of the Ct region (Supplementary Fig. S1). Among these structures, the largest conformational variation occurs in the intracellular section of the pore-lining helices M2 - M4, categorized into two distinct states (Fig. 1a-c): the up-state, where the M4 helix is orientated towards the membrane bilayer, resulting in closure of the intramembrane-facing side fenestrations and the down-state, characterized by M4 orientating away from the inner membrane leaflet with opening the side fenestrations. Due to its larger occupation of membrane volume, the up-state is thought to represent the highly active state induced by membrane stretching. The down-state is characterize be a high sensitivity to the inhibitors fluoxetine and norfluoxetine (NFx) that bind in the side fenestration and is thought to represent a low activity state^14,19,20^,. Alternative concepts have been proposed to explain the differences in activity between the up- and down-states. One concept, derived from the TRAAK crystal structures, suggests a lipid block mechanism^13^. Here, the two crystallographic states show identical and apparently conductive SF. In the down-state, however, an additional electron density below the SF is interpreted as the acyl-chain of a phospholipid, which could potentially block ion permeation. A very recently cryo-electron microscopy (cryo-EM) structure of TREK-1, obtained in the presence of the *n*-dodecyl-β-D-maltoside (DDM) detergent and phosphatidylethanolamine (PE), reports contrasting findings^21^. In the down-state, the positively charged head group of PE binds to the four threonine residues that make up the S4 K^+^ binding site in TREK-1, rather than an acyl-chain directly blocking the ion permeation pathway.

**Fig. 1.**
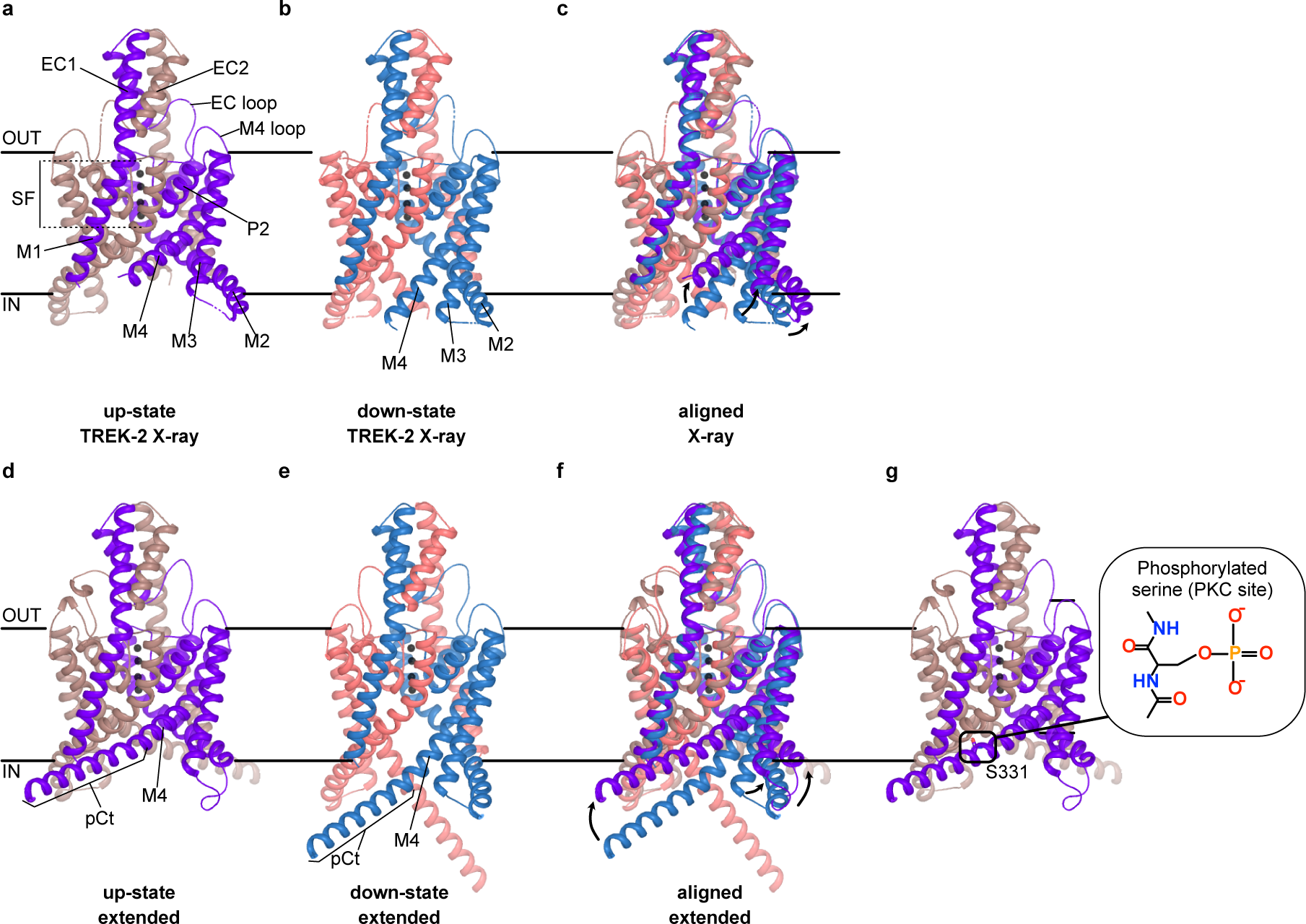
Structures and functional states of TREK-2 K_2P_ channels. **(a-c)** Crystal structures of the (a) up-state (PDB ID: 4BW5)^14^ and (b) down-state (PDB ID: 4XDJ)^14^ TREK-2 channel are shown, along with (c) an alignment of both structures. **(d-f)** The same structures depicted for TREK-2 with a 19-residue extension of the proximal C-terminus (pCt) in their helical conformation (designated as TREK-2*), which served as the starting structure for MD simulations. **(g)** TREK-2* structure as in (d) with S331 in the pCt highlighted. Note, S331 is phosphorylated in the respective MD simulations.

Another striking feature of K_2P_ channels is their ability to convert various external stimuli into electrical signals. TREK channels, in particular, can be efficiently gated by changes in transmembrane voltage, pH, temperature, mechanical force, polyunsaturated fatty acids like arachidonic acid (AA), bioactive lipids such as PIP_2_, and phosphorylation^22–26^, highlighting them as prototypic polymodal-regulated ion channels. Functional experiments revealed the SF as the main gate in TREK channels^27–30^, but structural changes in the SF have been only revealed at low potassium (K^+^) concentrations^16^. For sensing and integrating most physicochemical factors, the Ct has been suggested to play a critical role in channel gating. For instance, altering the Ct sequence in the TREK-2 renders it insensitive to pH and fatty acid^31^, while truncation of the Ct in TREK-1 channels results in the loss of activation AA and anesthetics^23,32^.

Moreover, TREK channel activity undergoes dynamic regulation by protein kinase A (PKA)-, C (PKC)- and G (PKG)-dependent signaling pathways that are thought to be involved in the development of hyperalgesia under inflammatory conditions and depression via the modulation of serotonin release in raphe nuclei^4,8,33,34^. Phosphorylation at two sites in the Ct has been revealed to be responsible for the inhibition downstream of receptor activation in TREK channels^33,35^. These two sites include S300 in TREK-1 (S331 in TREK-2) for PKC-induced phosphorylation and S333 in TREK-1 (S364 in TREK-2) for PKA-induced phosphorylation. Interestingly, it was shown that (de-)phosphorylation of TREK channels produces a dynamic interconversion between voltage-gated (phosphorylated) and ‘leak’-like (dephosphorylated) phenotypes^36^.

Direct structural evidence showing allosteric coupling between the proximal Ct (pCt) and the SF gate is currently lacking, as both up- and down-state TREK structures revealed the same SF conformation^14^, but several experimental and computational studies have suggested a possible coupling between different conformational states of the M4 helix and the SF gate^15,17,37–40^. However, previous molecular dynamics (MD) simulations were performed exclusively with the truncated form of TREK-2 lacking the pCt and consequently, the precise allosteric interaction network between these two dynamic hotspot regions remains poorly understood. To overcome this limitation, we incorporated a 19-residue pCt for both the up- and down-state structures of TREK-2 (referred to as TREK-2*; see method section) in our MD simulation and conducted an integrated investigation using functional and computational electrophysiology, encompassing (de-)phosphorylation mimics, state-dependent inhibitors, lipid tethering experiments, and extensive MD simulations. Our aim was to elucidate (i) the role of the pCt, (ii) pCt coupling to the SF, and finally (iii) the alterations in the filter gate defining TREK channel activity. This approach uncovered two distinct structural pathways by which up and down movements of the pCt controls the functional status of the SF being either conductive or non-conductive. The non-conductivity of the down-state resulted from a conformational change at the SF. Overall, our findings offer mechanistic insights into how different physiological signals like phosphorylation can be sensed and transmitted from the pCt to the SF in TREK K_2P_ channels at the atomistic scale.

## Results

### Modification of the pCt with alkyl-MTS reagents modulates TREK channel activity

To gain mechanistic insight into the regulation of TREK channel activity by the orientation of the pCt, we systematically introduced cysteine residues into the pCt of TREK-1 channels and attached probes of different chemical nature via methathiosulfonate (MTS)-mediated chemical modification. We hypothesized that introducing a lipophilic probe, such as an aliphatic chain (i.e. a decyl-chain), at sites facing the membrane would foster the interaction of the pCt with the membrane. Likewise, introducing lipophobic (hydrophilic) probes at these sites, such as charged moieties (e.g. MTS-ET^+^ or MTS-ES^-^), would destabilize membrane interactions. This approach aims to move the pCt towards or farther away from the membrane and, thereby allowing us to study the impact of this movement on TREK channel function.

First, we measured WT TREK-1, WT TRAAK and TREK-2* channels without modifiable cysteine residues as references. Upon application of 100 µM of decyl-MTS for at least 30 s to the respective channel in excised membrane patches no change in current amplitudes were recognized (Fig. 2a, Supplementary Fig. S2a,b). Note, patch integrity and channel regulation were tested with either 1 mM tetra-pentyl-ammonium (TPenA) and a short pH_i_ activation step. Next, we conducted systematic cysteine scanning mutagenesis of the entire pCt and examined the effect of charged and aliphatic MTS probes. We found that TREK channels carrying cysteine residues in the pCt in a regular interval could be activated robustly with decyl-MTS (Fig. 2a,b, Supplementary Fig. S2c). Remarkably, all hit residues face toward the inner leaflet of the lipid bilayer (Fig 2b, inlay). The strongest activation was observed for TREK-1 T318C, V322C and F325C, respectively, with full current increases of 33 ± 11-fold, 42 ± 17-fold and 103 ± 20-fold, respectively. Strikingly, the corresponding cysteine mutant channels in TRAAK and TREK-2 could be similarly activated with decyl-MTS (Fig.2 b,d,e). Furthermore, modification of the hit residues in TREK-1 (T318C, V322C and F325C) with charged MTS probes like MTS-ES^-^ and MTS-ET^+^ resulted in the opposite effect on activity, i.e. channel inhibition (Supplementary Fig. 2d). These results suggest a helical conformation for the pCt in the TREK channel family and that introducing lipid anchors mimicking decyl-groups to the membrane-facing site appears to lock all three TREK/TRAAK channels in a highly active state.

**Fig. 2.**
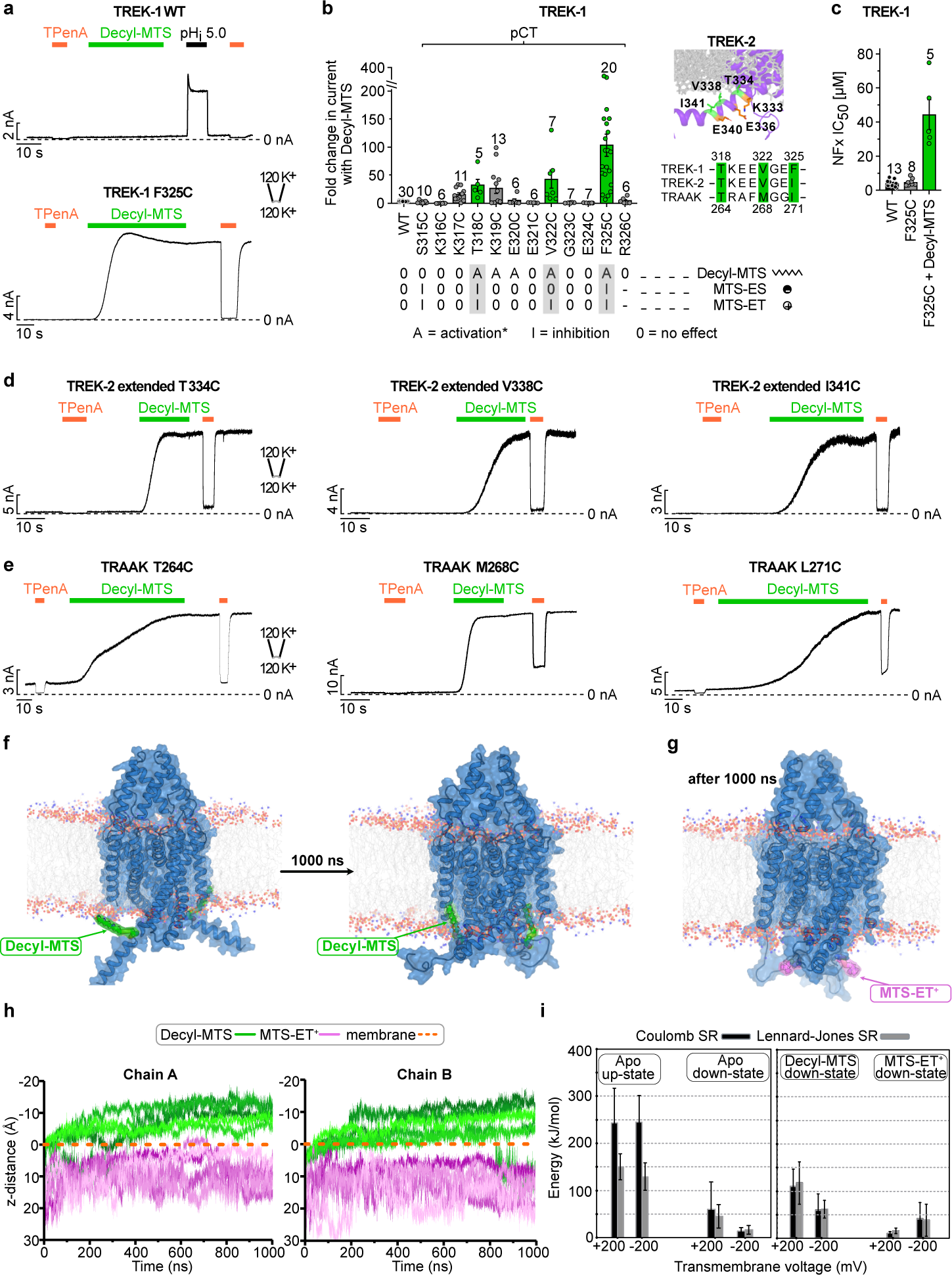
Membrane tethering of the pCt with cysteine modifying probes activates TREK channels. **(a)** Example recordings of WT and F325C mutant TREK-1 channels in a symmetric K^+^ gradient at pH 7.4 at +40 mV showing no effect upon application of 100 µM decyl-MTS for WT and robust activation for F325C mutant channels. **(b)** Systematic screening of cysteine mutants in the pCt region of TREK-1 by recordings as in (a) reveal mutants that are activated (A) with decyl-MTS. Note, side chains of mutant channels that are strongly activated (green sticks) point to the inner leaflet of the lipid bilayer phase (gray sticks) as highlighted for TREK-2* in the inlay. Mutant TREK-1 channels with robust decyl-MTS activation show an inhibition (I) upon MTS-ET^+^ application. **(c)** Apparent affinities (IC_50_ values) of NFx for WT, F325C mutant and decyl-MTS-activated F325C mutant TREK-1 channels. **(d,e)** Example recordings as in (a) of TREK-1 homologous mutant channels with the strongest decyl-MTS response in TREK-2* (d) and TRAAK (e). Note, TREK-2* for functional recordings corresponds to the construct used in MD. **(f)** Representative initial (left) and final (right) snapshots of decyl-MTS-modified TREK-2* from a 1 μs MD simulation. (**g**) A representative final snapshot of MTS-ET^+^-modified TREK-2* from a 1 μs simulation. **(h)** The distance (D_z_ component of the vector) between a chosen atom of the ligand (C12 for decyl-MTS, N2 for MTS-ET^+^) and the membrane plotted against simulation time under +200 mV. The membrane was defined as a plane by selecting the center of geometry of phosphorus atoms located in the cytosolic part of the POPC lipid bilayer. **(i)** pCt-membrane interaction energies derived from simulations of apo TREK-2* (up- and down-state) as well as MTS-modified TREK-2* starting from the down-state structure under −200 mV and +200 mV transmembrane voltage. Averaged short-range Coulomb (Coulomb SR) and short-range Lennard-Jones (Lennard-Jones SR) energies were calculated. Values are given as mean ± s.e.m with number (n) of experiments indicated in the figure.

We further investigated this structural transition using MD simulations and *in silico* attaching either decyl-MTS or MTS-ET^+^ to the cysteine at position 334 (T334C) of TREK-2* (corresponding to TREK-1 T318). For both decyl-MTS- and MTS-ET^+^-bound channels, we conducted five runs of 1 μs simulations, starting from the down-state TREK-2* structure. As expected, in the decyl-MTS-modified channel, we observed the insertion of the decyl-chain chain into the lipid bilayer (Fig. 2f), while membrane embedding of the -ET^+^ moiety was never observed (Fig. 2g). This differential behavior was supported by calculating the distance between the ligands and the lipid bilayer in the z-direction (Fig. 2h, Supplementary Fig. S3). These results suggest that attaching a decyl-group to the pCt promotes movement towards the membrane. Whether this induces a state similar to the crystallographic up-state was further tested functionally by using the state-sensitive inhibitor NFx, as NFx only binds to the down-state. To this end we measured NFx inhibition for the TREK-1 F325C mutant channels before and after decyl-MTS modification. Before modification, F325C mutant channels showed a similar high sensitivity to NFx as WT channels, indicating a large fraction of the channels in the NFx-sensitive down-state. In contrast, decyl-MTS modification caused a dramatic drop in NFx sensitivity (Fig. 3c) indicating that a large fraction of channels now exists in an up-state like conformation.

**Fig. 3.**
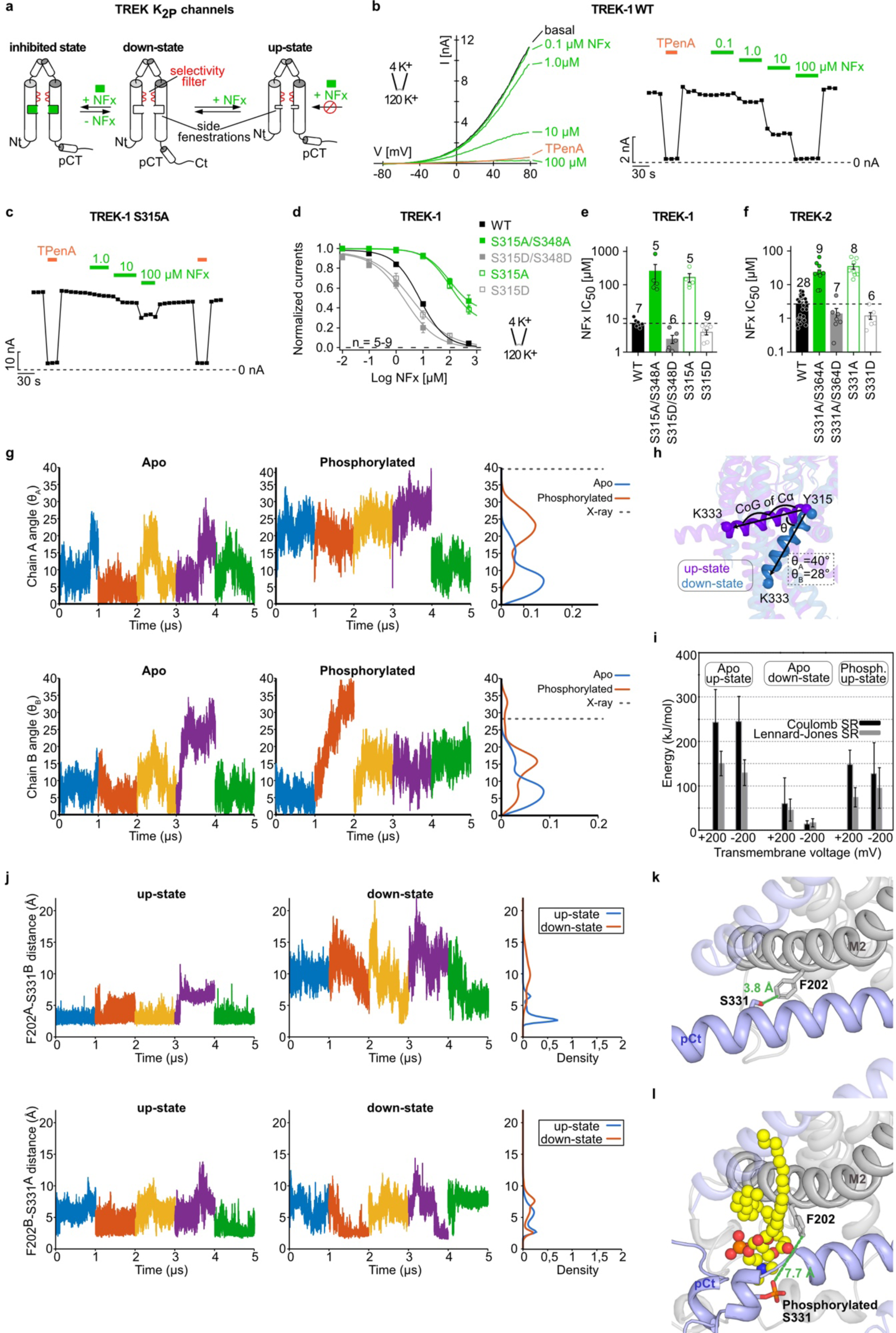
Phosphorylation-mediated state transition of TREK channels. **(a)** Cartoon depicting gating model and state dependency of NFx inhibition in TREK K_2P_ channels. **(b)** Example recording of WT TREK-1 channels in an asymmetric K^+^ gradient at pH 7.4 in a voltage range from −80 to +80 mV showing the dose-dependent inhibition with the indicated concentrations NFx (left) and the time course of block analyzed at +40 mV (right). **(c)** Same recording and analysis as in (b) for S315A TREK-1 channels. **(d)** Dose-response curves of NFx inhibition for WT and mutant TREK-1 channels as indicated analyzed from recordings as in (b). **(e,f)** NFx IC_50_ values from dose-response curves as in (d) for WT and mutant TREK-1 (e) and TREK-2 (f) K_2P_ channels. **(g)** Time course of the θ-angle for M4 in the apo and phosphorylated TREK-2* simulations starting from the up-state structure, shown for the first and second chains. Histograms of θ-angle distribution are included on the right site. θ-angle of the up-state and down-state X-ray structures was used as a reference (**h**) and shown as dashed line. **(i)** pCt-membrane interaction energies derived from simulations of apo TREK-2* (up- and down-state) as well as phosphorylated TREK-2* starting from the up-state structure under −200 mV and +200 mV. Averaged short-range Coulomb (Coulomb SR) and short-range Lennard-Jones (Lennard-Jones SR) energies were calculated between the pCt and membrane. Values are given as mean ± s.e.m with number (n) of experiments indicated in the figure. **(j)** Time course of the inter-chain distance between S331 and F202 for both up- and down-state simulations of TREK-2*. Histograms of distance distribution are included on the right site. **(k)** The distance between S331 and F202 was illustrated for the unphosphorylated TREK-2* in the up-state and **(l)** for a phosphorylated TREK-2* after 1 μs simulation.

### Phosphorylation at the PKC site in the pCt promotes the up-to-down transition

Having established that up and down movement of the pCt controls TREK channel activity, we explored whether a physiologically relevant stimulus employs the same mechanism. We focused here on the regulation of TREK channels by PKC and PKA, as both enzymes exert a strong inhibitory effect via phosphorylation sites at the Ct. The PKC site is located in the pCt (S300 in mTREK-1 and S326 in rTREK-2), whereas the PKA site is located at a more distal site in the Ct (S333 in mTREK-1 and S359 in rTREK-2)^33,35,41^. Recently, it was shown that in the process of receptor- and kinase-mediated modulation of TREK channels a sequential phosphorylation is responsible for the inhibitory effect on channel activity^33^ with PKA phosphorylation promoting PKC phosphorylation. In the same study it was shown that alanine substitutions at both sites (S300A/S333A) mimicking a permanently dephosphorylated state enhance TREK-1 channel currents, while a double aspartate mutation (S300D/S333D) mimicking permanently phosphorylated channels drastically reduced TREK currents.

To investigate the mechanistic basis for phosphorylation regulation of TREK channels we first tested if the (de-)phosphorylated states with enhanced or diminished activity can be assigned to either the up- or down-state conformation of TREK utilizing as before the state-sensitive inhibitor NFx (Fig. 3a)^14,20,42^. We measured TREK-1 channel currents in response to 800 ms voltage ramps (−80 mV to +80 mV) in excised membrane patches under asymmetrical K^+^ conditions (Fig. 3b). TREK-1 WT is inhibited by NFx with an apparent affinity of ∼7 µM (IC_50_ = 7 ± 1 µM, n = 7) (Fig. 3b,d,e). In agreement with previous studies, the double alanine substitution in hTREK-1 (S315A/S348A; equivalent to S300A/S333A) and hTREK-2 (S331A/S364A; equivalent to S326A/S359A) resulted in largely increased currents (Supplementary Fig. S4a). Notably, NFx sensitivity of these currents were strongly reduced and even high concentrations produced incomplete inhibition (IC_50_ = 249 ± 149 µM, n = 5 for TREK-1 and IC_50_ = 24 ± 4 µM, n = 9 for TREK-2) (Fig. 3d-f, Supplementary Fig. S4b). In contrast, the phosphorylation mimicking substitutions S315D/S348D in TREK-1 and S331D/S364D in TREK-2, respectively, resulted in smaller currents (Supplementary Fig. S4a) and slightly increased NFx sensitivities compared to WT (IC_50_ = 2 ± 1 µM, n = 6 for TREK-1 and IC_50_ = 1 ± 1 µM, n = 7 for TREK-2) (Fig. 3d-f, Supplementary Fig. S4c). Further, the PKC site (S315) appeared to dominated the overall impact of phosphorylation on TREK activity as the (de-)phosphorylation mimicking alanine (S315A) or aspartate (S315D) substitutions in TREK-1 and TREK-2 had a similar impact on NFx sensitivity and current amplitude as seen for the respective double substitutions (Fig. 3c-f, Supplementary Fig. S4d,e).

These results indicated that mutations at the PKC site mimicking dephosphorylation induce a down-to-up state transition in TREK channels, while phosphorylation mimics retain the channels in a down-state conformation. This also implies that TREK channels at least in our expression system are strongly phosphorylated without external kinase stimulation as introducing the phosphomimic substitutions only slightly increase the NFx sensitivity.

We further explored the impact of phosphorylation at the PKC site with MD simulations and adding a phosphate group *in silico* at residue S331 of TREK-2* in the up-state (Fig. 1g). We performed five runs of 1 μs MD simulations under either −200 mV or +200 mV transmembrane voltages to assess if phosphorylation would trigger a shift from the up- to the down-state. We focused on the movement of the M4 helix and used the θ-angle that represents the angle difference of the M4 helix in the up-state in respect to the down state as readout for the state transition (Fig. 3g). Crystal structures showed an angle range of 28 - 40° between M4 in the up- and down-states. In the apo-TREK-2* simulations starting from the up-state, the θ-angle remained mostly small, indicating no obvious conformational transition. However, in the phosphorylated simulations, we observed a notable shift towards larger θ distribution (Fig. 3g,h). While the θ-angle did not reach the full angle increase seen in the up- to down-state X-ray structure transition, it clearly indicates a trend toward the transition from the up-state to the down-state within the current time scale of the simulations. Importantly, our simulations revealed that phosphorylation at S331 disrupts the hydrophobic interaction between S331 at the pCt and F202 at M2 from the opposite subunit (Fig. 3j-l). Following the reduction of this hydrophobic interaction, we observed in some of the simulation runs that a lipid occupied a site between M2 and M4 (Fig. 3l). As expected, calculations of the short-range Coulomb and short-range Lennard-Jones energies between the pCt and the membrane revealed a significant energy decrease when phosphorylation was introduced during the up-state simulations (Fig. 3i). Collectively, our functional and computational data suggest that phosphorylation at the PKC site promoted a downward movement of the pCt leading to a conformational state resembling the down-state in the X-ray structure.

### The difference in ion conductivity is a result of the conformational change in the SF of TREK-2*

Our findings so far have established that the upward movement of the pCt induces a highly active channel state that likely resembles the crystallographic up-state. Previously, we and other have shown that TREK channel activity is controlled by a gate residing in the SF, but the crystallographic up- and down-states of TREK have identical SF structures, which argues against the contribution of the SF in TREK channel gating. However, the crystallographic conditions could have altered the energetics of the SF, thereby obscuring its intrinsic properties. Therefore, we performed MD simulations of K^+^ permeation in up- and down-state of TREK-2 with an extended pCt (TREK-2*). All simulation runs of the up-state TREK-2* were conductive at positive voltages (+200 mV) throughout the entire 1 μs MD runs (Fig. 4a, Supplementary Table S1). In a typical 1 μs trajectory under +200 mV, we observed an average of five K^+^ permeation events, corresponding to a conductance of 4 pS. Under negative voltage (−200 mV), four out of five simulations showed conductivity, albeit with a significantly lower conductance compared to the positive ones. In strong contrast to the up-state, no K^+^ conduction was observed in the down-state TREK-2* simulations (Fig. 4a, Supplementary Table S1). Therefore, our results reveal a strong correlation between ion conductivity and conformational state of pCt/M4. Similar to the previous simulations of down-state TREK-2 channel by Aryal *et al.*^43^, we also observed lipid penetration from the lateral side fenestration during the simulations. This took place mostly in the down-state simulations, but was also seen in some of the up-state runs (Supplementary Fig. S5) and did not obviously block ion conduction.

**Fig. 4.**
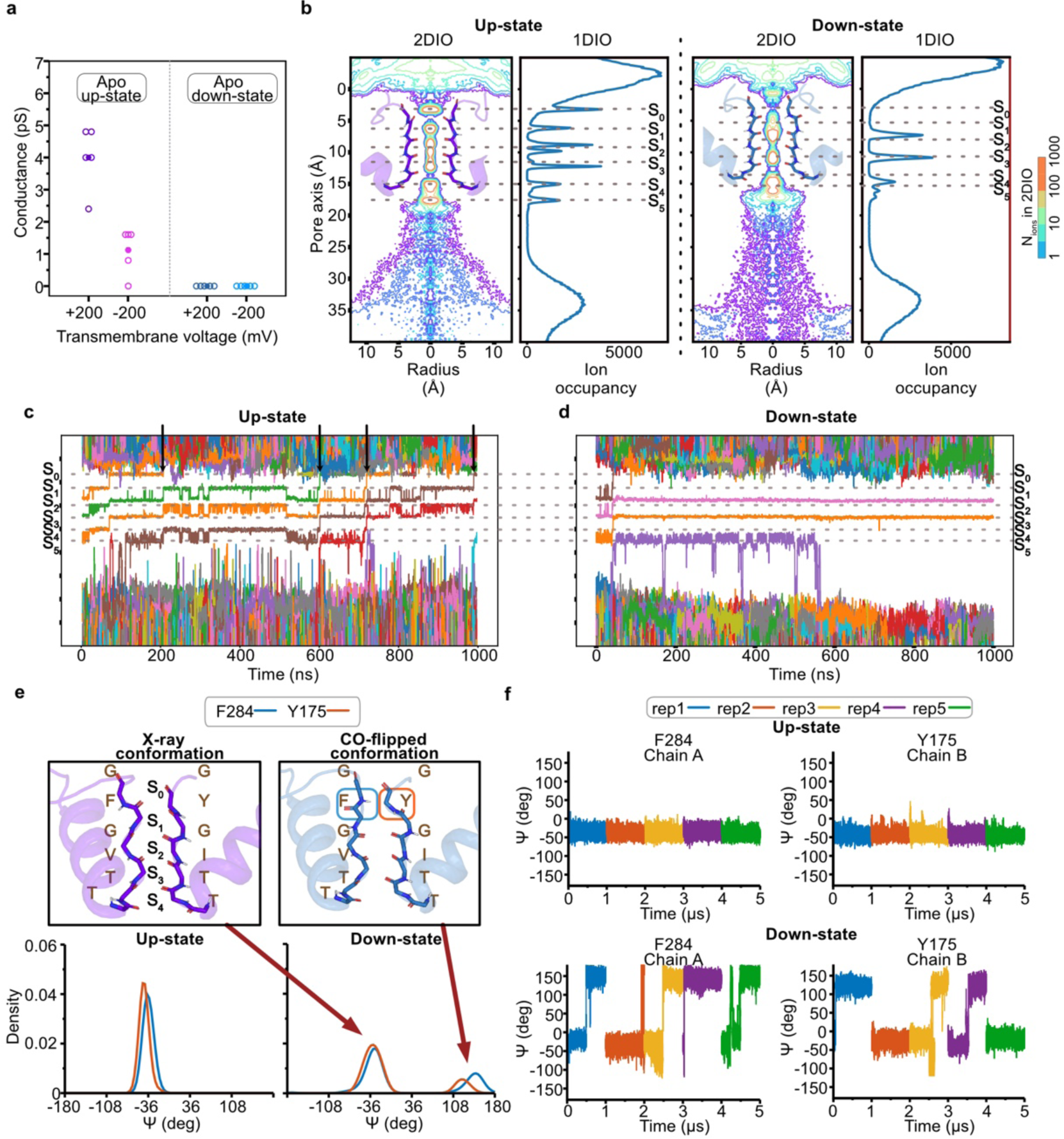
Conductance, SF ion occupancy and conformational changes determined from TREK-2* simulations. (**a**) Ion conductance derived from apo TREK-2* simulations in its up- (PDB ID: 4BW5)^14^ and down-state (PDB ID: 4XDJ)^14^. Filled and empty circles represent mean and individual ion conductance, respectively. (**b**) One- and two-dimensional ion occupancy profiles within the SF for combined up-state and non-conductive down-state TREK-2* simulations. The radial area of the pore was defined and ions passing along the pore axis (D_z_) were calculated from the simulations. The occupancy of ions was normalized per 0.001 Å^3^ per 1 μs based on the volume change along the radius. The center of mass of the SF backbone atoms was located on the 7 Å point along the pore axis. Ion binding sites along the pore axis were indicated with dashed lines. (**c**) Traces of ions passing through the SF during a representative 1 μs up-state and (**d**) down-state TREK-2* simulation. (**e**) Distribution of Psi angle (ψ) for F284 and Y175 from the combined up-state (left) and down-state (right) simulations. (**f**) Psi angles (ψ) variations of F284 in chain A and Y175 in chain B over time in the up-state (top) and down-state (bottom) of TREK-2*. Individual simulation runs were carried out with AMBER99sb^44^ for 1 μs under a transmembrane voltage around +200 mV.

Analysis of one- and two-dimensional ion occupancies suggested significant differences in the preferred K^+^ binding sites in the SF region between the up- and down-state (Fig. 4b, Fig. S6) simulations of TREK-2*. While ion binding at sites S0 - S5 was almost equally populated during the conductive up-state simulations, ion binding at S1 and S4 was abolished in the simulations of down-state TREK-2 (Fig. 4c/d). Due to significant differences in ion occupancy between up- and down-state simulations, we further compared the SF backbone conformation in these simulations (Supplementary Fig. S7). Remarkable differences were identified for Y175 and F284 around the S1 K^+^ site (Fig. 4e). While the crystallographic SF conformation remained stable throughout the entire conductive up-state runs, in the non-conductive down-state simulations, the backbone carbonyl of Y175 and F284 flipped away from the ion conduction pathway in most trajectories and remained stable for the remainder of the simulation time (Fig. 4e,f, Supplementary Fig. S8-9). Building upon previous MD studies that demonstrated carbonyl flipping in the SF can impede ion conduction in K^+^ channels^16,39^, we propose that the ion non-conductivity in the down-state simulations resulted from the carbonyl flipping of Y175 and F284 in TREK-2*, leading to a strong reduction in the ion occupancy at S1 and S4.

### Interaction network analysis revealed the coupling mechanism between SF gate and pCt in TREK-2*

To gain further insights into the precise energetic coupling between the pCt/M4 and the SF, we conducted an extensive interaction network analysis. We identified intra- and inter-chain contacts between residues within a 5 Å radius during up- and down-state simulations, respectively. By subtracting the population of down-state contacts from the up-state ones, we constructed a contact difference map to highlight key disparities. In total, we identified 86 contact differences (> 0.25 probability), comprising 22 inter-chain (transmembrane helices of both channel subunits) contacts and 64 intra-chain (transmembrane helices of the same channel subunit) interactions (Table S2).

Analyzing the contact differences, first of all we observed substantial variations between up- and down-states at the interfaces of M2 and pCt/M4 (Fig. 5). Specifically, we identified five hydrophobic contacts predominantly present in up-state simulations between the pCt/M4 and M2 helices from opposing subunits. One of these involved P198 from the M2 helix, engaged in three of the five identified contacts (with F316, L320 and I323 from M4). S331 - F202 and F316 - I197 constituted two other key inter-chain contacts distinguishing the up- and down-states. In contrast, the key inter-chain interactions between M4 and M2 were largely disrupted in the down-state simulations, being replaced by new interactions between M4, M2 and M3 within the same subunit, such as W326 - R237, R237 - E223 and R237 - K224 (Fig. 5a). We computed the distance distribution of the inter-chain interaction P198 - L320 and intra-chain interaction L211 - A318 throughout the simulations. In the up-state simulations, the distance between P198 and L320 remained small, even slightly smaller compared to the crystallographic state in one subunit (Supplementary Fig. S10a). However, in the down-state simulations, this interaction became highly asymmetric and was exclusively abolished in one subunit. This trend was inverted for the intra-chain interaction L211 - A318 (Supplementary Fig. S10b). Notably, this result is in full consistency with the finding of the phosphorylated TREK-2* simulations, where the disruption of the hydrophobic interaction between S331 and F202 was found to be critical for inducing the down-state.

**Fig. 5.**
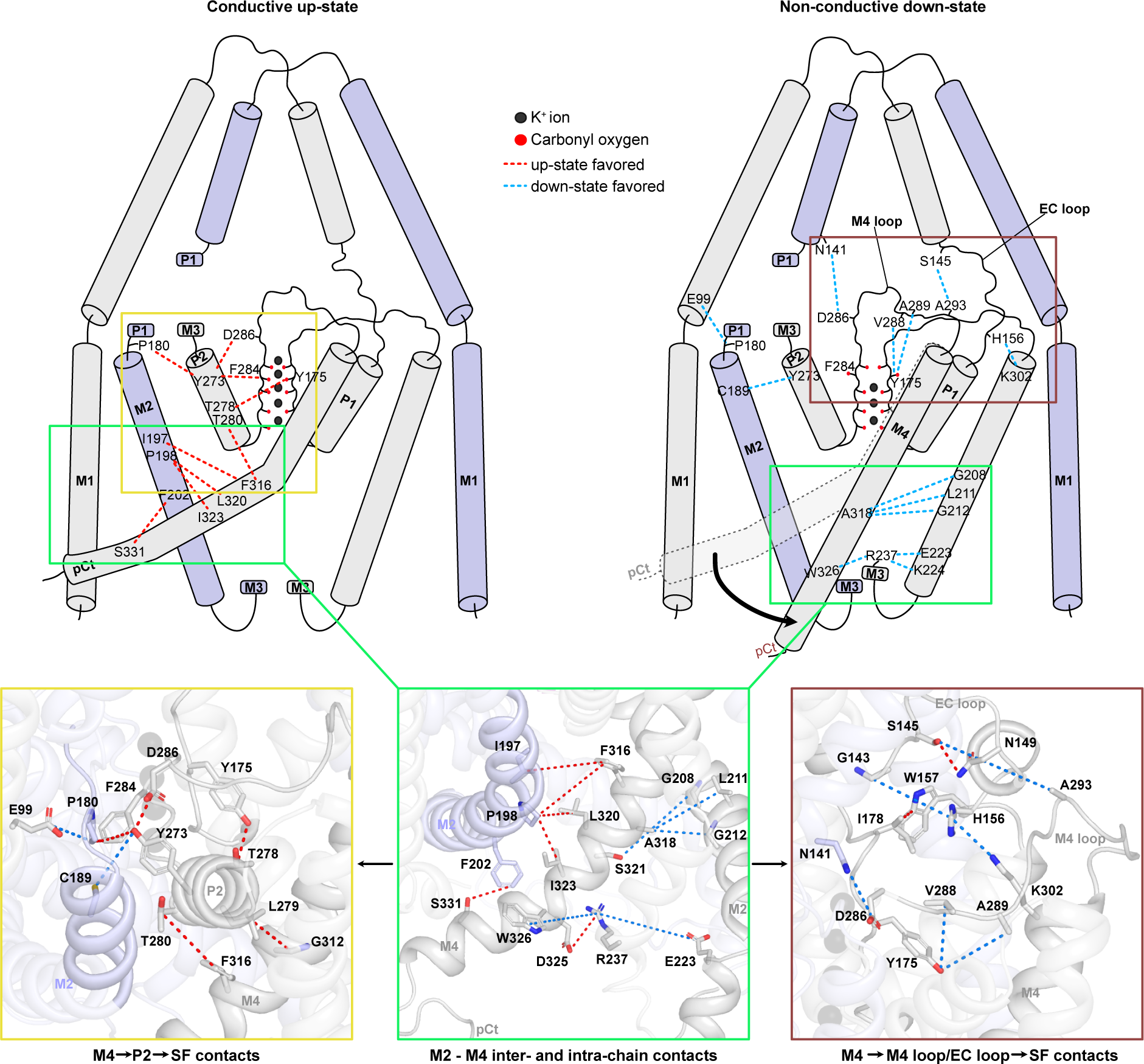
Coupling mechanism between pCt/M4 and SF gate. pCt/M4 dynamics is coupled with the M2 - M4 inter- and intra-chain interactions that further coupled to the SF gate via two different pathways: (i) M4 → P2 → SF; (ii) M4 → M4 loop/EC loop → SF. Favorable interactions in the up-state are depicted with red dashed lines, while blue dashed lines represent favorable interactions in the down-state. Bottom middle: key M2 - M4 inter- and inter-chain contacts. Bottom left: the first pathway of the pCt/M4 → SF coupling involves a number of M4 → P2 → SF contacts. Bottom right: the second pathway of the pCt/M4 → SF coupling involves the M4 → M4 loop/EC loop → SF contacts.

Triggered by the differences in interactions between M2, M3 and M4, the dynamics of the pCt/M4 and the SF were further coupled through two distinct pathways. The first pathway involves the M2 helix and the P2 pore-forming domain (Fig. 5, primarily inter-chain interactions), where interactions such as I197 (M2) - F316 (M4), F316 (M4) - T280 (P2), Y273 (P2) - F284 (SF2), T278 (P2) - Y175 (SF1), Y273 (P2) and P180 (P1-M2 linker) were predominantly observed in the up-state simulations. Conversely, in the non-conductive down-state simulations, a series of new interactions emerged, including P180 (M2) - E99 (M1) and Y273 (P2) - C189 (M2). In the second pathway, coupling of the pCt/M4 and the SF was mediated by the M4 loop, which connects SF2 and the upper M4 helix (Fig. 5). Several differences were identified between the M4 loop and its interacting counterparts. For instance, the interaction between Y175 (SF1) and T278 (P2) stabilized the conductive SF conformation in the up-state simulations, but this interaction diminished and was replaced by the Y175 (SF1) - V288 and A289 (M4 loop) interactions. Another example is the dominant interaction between Y273 (P2) and D286 (M4 loop) in the up-state, which was replaced by the D286 (M4 loop) - N141 (EC2) interaction in the down-state simulations. Overall, these key interaction differences in interactions identified in these two pathways provide valuable atomistic insights into how the dynamics of the pCt/M4 and the SF are coupled to each other.

## Discussion

TREK K_2P_ channels constitute background K^+^ channels that are susceptible to regulation via G-protein-mediated pathways and protein kinases^33,35^. Phosphorylation at the Ct by PKG, PKA and PKC inhibits TREK channels. These kinases target specific Ct phosphorylation sites, with cross-influence on each other’s activity. Notably, PKC possesses a phosphorylation site in the pCt domain. Previous studies have also suggested the essential involvement of the pCt in PIP_2_ activation^45^, pH regulation, and mechanosensitivity^46^. In this study, based on electrophysiological measurements of (de-)phosphorylation mimics at the PKC site, we were able to demonstrate that phosphorylation strongly affects the up- and down-state conformational equilibrium. These results have been further substantiated by atomistic MD simulations, revealing a clear trend of conformational transition from the up- to down-state upon phosphorylation at the PKC site and reduced pCt-membrane interaction.

Modifying the pCt with alkyl-MTS probes of varying hydrophobicity yielded contrasting channel activities and conformational equilibria between up- and down-state, further confirming the crucial regulatory role of the pCt domain in TREK channels. While the interaction of the pCt/M4 and the membrane was reinforced by embedding decyl-MTS into the membrane, MD simulations of the MTS-ET^+^ variant did not demonstrate the same side-chain insertion, leading to a reduced interaction between pCt and membrane. In decyl-MTS-modified simulations, we did not observe a full down-to-up transition, most likely due to the limited time duration of the simulation timescale.

Microsecond MD simulations of apo up- and down-state TREK-2 with extended pCt revealed a striking difference in the conductance of two distinct conformational states, particularly regarding the outward flow of K^+^. Debates still persist regarding whether the less-conductive down-state arises from a lipid pore block^13^ or conformational changes in the gating region. While we observed lipid penetration from the lateral side fenestration in several down-state TREK-2* simulations, similar to previous simulations^43,47^, we also observed the same in some of our up-state simulations without significant obstruction of channel conductance (Supplementary Fig. S5).

MD simulations revealed significant differences in S1 K^+^ occupancy between up- and down-state TREK-2*. Non-conductive down-state simulations showed reduced S1 ion density due to the reorientation of the backbone carbonyls of Y175 and F284. Although no conformational difference was observed in the SF of the X-ray structures between the up- and down-state TREK-2, we noticed that ion density at the S1 site was only present in the up-state structure^14^. The absence of ion occupation at the S1 site in the down-state X-ray structure reflects the dynamic nature of this ion binding site. In line with that, another structural study revealed substantial conformational changes relayed to the S1 and S2 K^+^ sites in the X-ray structures of TREK-1 at low K^+^ concentrations^16^. Similarly, cryo-electron microscopy (cryo-EM) structures of the non-conductive state of TASK-2 K_2P_ channels showed conformational changes of F101 in SF1 and F206 in SF2 at the S1 ion binding site.^48^ These observations suggest a consensus mechanism for C-type gating in the SF within the K_2P_ channel family, primarily focused on the S1 site. In fact, a comparison of the SF sequences between most K_2P_ channels (except TWIK channels) and canonical K^+^ channels revealed a major difference: one (in SF2) or both (in SF1 and SF2) tyrosine (Tyr) residues, adjacent to the S1 K^+^ site, were replaced by phenylalanine (Phe) (Supplementary Fig. S11). This sequence alteration leads to a reduction in hydrogen bonding interactions between the SF and the pore-forming helix, resulting in decreased SF stability. hERG (human Ether-a-go-go Related Gene) channels, also with a Phe at the homolog position in the filter, exhibits side-chain shifts in the cryo-EM structures that were suggested to be associated with fast inactivation^49^. Nevertheless, we propose that the configuration without ions at the S1 site, coupled with the carbonyl reorientation of Y175 and F284, does not represent the true SF conformation of inactivated TREK-2* channels. Previously, from the gating charge analysis, we proposed the inactive filter to be in an ‘ion-deplet’ state^27^. Therefore, the depletion of ions at the S1 site in the SF may only represent the initial step of TREK-2 inactivation, while the complete inactivation process could not be elucidated in the current study due to simulation time limitation.

Based on the functional and simulation data discussed above, a strong connection between the pCt/M4 dynamics and C-type gating in the SF has been established. The mechanism is illustrated in Fig. 6.

**Fig. 6.**
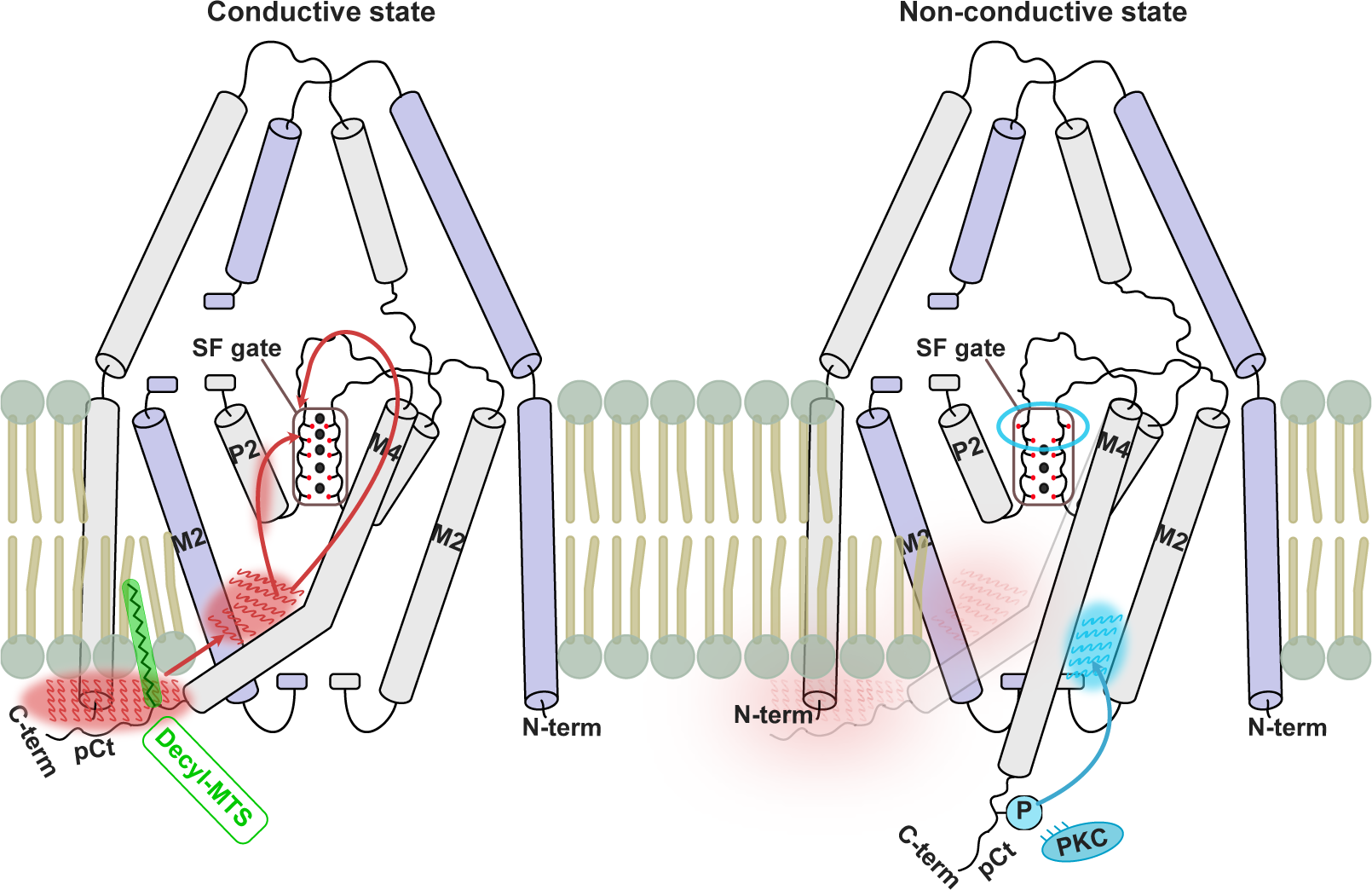
Schematic illustration of TREK-2 K_2P_ channel gating and modulation. The conductive state is stabilized by the strong interaction between pCt and the membrane, promoting inter-chain M4 - M2 interactions. These interactions are coupled with the SF gate through two pathways, thereby stabilizing the conductive state of the SF (left). Disruption of the interactions between pCt and the lipid bilayer leads to the emergence of new interactions between the M4 and M2 of the same subunit, resulting in conformational changes of the SF that leads the channel to a non-conductive state (right). Decyl-MTS (green) enhances the interactions between pCt and the membrane. In contrast, phosphorylation of S331 at the pCt (blue) destabilizes the interactions between pCt and the membrane, leading to a non-conductive state of the channel.

Firstly, substantial variations in the M4 and M2 interaction interfaces were identified between the up- and down-states. In the conductive up-state simulations, inter-chain interactions between M4 and M2 dominated, whereas in the non-conductive down-state simulations, this simulation network was diminished and replaced by intra-chain interactions of M4, M2 and M3. Phosphorylation at S331 disrupts the hydrophobic interaction between pCt/M4 and M2 from the opposite subunits, thereby promoting the down-state. Triggered by these differences in the interaction network, two key interaction pathways, that mediate the coupling between M4 and the SF have been further identified: (i) M4 → M4 loop/EC loop → SF, where interactions between the M4 - loop and surrounding residues (e. g. in EC loops) facilitate the conformational transition in the upper part of the SF. This finding aligns well with previous functional and simulation studies, which suggested that M4 - loop dynamics play a crucial role in C-type gating^16^. (ii) M4/M2 → P2 → SF, where the cross-talk between M4 dynamics and the SF gate is mediated by the interactions between M2, M4, and P2 (Fig. 5). Particularly for the TREK-2*, a strong hydrogen bonding between the side chains of Y175 (SF1) and T278 (P2) was observed in the up-state simulations, but this interaction was partially diminished in the down-state one. This result is in line with several recent solid-state NMR studies that emphasize the vital roles of the interactions between the SF and the pore helix in stabilizing the conductive SF conformation of K^+^ channels^50,51^. Moreover, the interaction between the P2 and M4 helix is primarily mediated by the interaction between F316 (M4) and T280 (P2), an interaction that was predominant only in the up-state simulations. Interestingly, the significance of this interaction in mediating the coupling between the M4 helix and the SF had already been proposed in previous simulations of the K_2P_ channel TRAAK^52^.

In conclusion, using an integrated approach, combining functional and computational electrophysiology, including systematic cysteine scanning mutagenesis, lipid tethering and blocker experiments, molecular dynamics simulations and interaction network analysis, we have firmly established the essential role of the pCt in TREK channel gating. Furthermore, these finding have facilitated a comprehensive understanding of how the interaction between the pCt domain and the membrane influence the conformational equilibrium between the up- and down-state of TREK channels, which is further coupled with the SF gate. By conducting extensive MD simulations, we have identified the conformational changes in residues near the S1 K^+^ site in the filter of TREK-2 that contribute to the initial step of C-type inactivation. This result aligns with numerous prior structural studies, shedding light on the dynamic nature of the S1 K^+^ binding site. Importantly, we have proposed two precise pathways that delineate the energetic coupling between the cytosolic sensing domain and the SF. These pathways are consistent with previous functional and mutational studies, providing a comprehensive understanding of how the physiologically highly relevant phosphorylation process leads to channel inhibition at the atomistic scale.

## Methods

### Computational electrophysiology

We started MD simulations from the high-resolution X-ray structure of up- (PDB ID: 4BW5)^14^ and down-state (PDB ID: 4XDJ)^14^ TREK-2 channels, respectively. The missing pCt in each subunit was manually extended using the PyMOL builder option^53^. To preserve the helical conformation, the helix option was selected during the addition of the amino acid sequence. The pCt was extended by adding 19 residues (^333^TKEEVGEIKAHAAEWKANV) based on the TREK-2 construct^54^. We added N-methyl amide and acetyl caps to the N and C termini using the PyMOL builder option. SWISS-MODEL^55^ was used to build missing loops (residues 149 to 154 and 229 to 235).

We performed MD simulations with AMBER99sb^44^ force field. The insertion of the TREK-2 channel into a POPC was performed with the GROMACS internal embedding function. The concentration of KCl was 600 mM in simulations. Improved ion parameters^56^ and lipid parameters^57^ were employed in the simulations. The SPC/E water model^58^ was used in simulations.

All MD simulations were performed with the GROMACS software (2019.3 and 2021.2)^59^. Short-ranged electrostatics and van der Waals interactions were truncated at 1.0 nm. Long-range electrostatics were calculated with the Particle-Mesh Ewald summation^60^. Temperature and pressure coupling were treated with the V-rescale^61^ scheme and the Parrinello-Rahman barostat^62^, respectively. The temperature was set to 300 K and the pressure to 1 bar. Fluctuations of the periodic cell were only allowed in the z-direction, normal to the membrane surface, keeping the density of the membrane unchanged. Verlet cut-off scheme was used for neighbor searching and pair interactions^63^. All bonds were constrained with the Linear Constraint Solver (LINCS) algorithm^64^. We used virtual sites for hydrogens to decrease the computational cost of apo- and phosphorylated TREK-2 simulations. These simulations were performed with an integration time step of 4 fs. All other TREK-2 simulations with covalently bound MTS reagents used an integration time step of 2 fs.

For the in-silico attachment of the MTS reagents, methyl sulfonyl groups of decyl-MTS and MTS-ET^+^ were removed to represent solely the side-chain structure, respectively. A cysteine residue was fused by a disulfide bridge to the trimmed reagent. Cysteine residue was capped with N-methyl amide and acetyl capping groups. The geometry of the structure was optimized and the electrostatic potential was calculated at the Hartree-Fock/6-31G* level using Gaussian16^65^. The generalized amber force field (GAFF)^66^ topology of the structure was generated with the Antechamber software^67^ using partial charges from the preceding quantum mechanics calculations according to the restrained electrostatic potential (RESP) approach^68^. After the geometry optimization, caps and cysteine were removed from the structure. The remaining side chain was fused by a disulfide bridge to the T334C mutation located in pCt. The disulfide bridges were generated at a distance of 2.05 Å.

After the channel was embedded into POPC with ions, the system was energy minimized and equilibrated. The energy minimization was done with GROMACS “steepest descent” algorithm with a maximum energy value of 100 kJ mol^-1^ to make sure to minimize the energy of the system as much as possible. After the minimization, the system was equilibrated for 10 ns with a position restraint by a force constant of 1000 kJ mol^-1^ nm^-2^ in x,y, and z-direction on the backbone atoms of the protein followed by 20 ns without position restraints on the backbone atoms of the protein. During the complete equilibration, the isothermal–isobaric (NPT) ensemble was used.

A copy of the equilibrated TREK-2 system was prepared and two identical copies were stacked on top of each other in the z-direction in order to construct a CompEL setup. Another energy minimization was performed for the new double-bilayer system to prevent crashes of molecules located in the intersection point of the two systems. An ion difference of 2 was introduced between the two compartments separated by lipid bilayers to introduce a transmembrane potential of about ±200 mV. Additional particle interchange algorithm (deterministic protocol) were employed during MD simulations to keep the number of ions in every compartment constant alchemically^69^.

All trajectories were analyzed with GROMACS tools and Python using MDAnalysis^70^ together with NumPy^71^ matplotlib^72^ and SciPy^73^. Distances were calculated with GROMACS tool *gmx dist* and dihedral angles were calculated with *gmx rama*. Ion occupancies and the trace of ions were estimated with Python using MDAnalysis by aligning different simulation setups using the center of mass of the backbone atoms of the SF residues as a reference point. Presented data of ion occupancies were derived from the combined trajectories of five independent replicates from +200 mV in the up-state (Fig. 4b), as well as five independent replicates from +200 mV in the down-state. Presented data of dihedral angle distributions were calculated using the kernel density estimation of combined trajectories of five independent replicates from +200 mV (Fig. 4e).

To calculate the θ-angle of the M4 helix, we designated the Cα atom of the Y315 residue as the initial point of the vector. The second point of the vector was defined as the geometric center of the Cα atoms of residues located between Y315 and K333. Using a combination of MDAnalysis and NumPy, we computed the vector difference of M4 for each frame relative to the reference X-ray structure.

The interaction energy between pCt and the membrane was determined using the GROMACS software tool, specifically the *gmx energy* module. The calculation of the interaction energy involved considering pCt starting from the residue S331. The membrane was represented by selecting all the atoms of the POPC molecule. To ensure statistical significance, five independent replicates were performed to calculate the mean and confidence interval of the interaction energy.

The interaction network analysis was performed using the combined trajectories of up-state apo TREK-2 at +200mV, each comprising 1 μs simulation run. The same was repeated for the combined trajectories of non-conductive down-state TREK-2 trajectories. A pair of residues were considered to be in contact if the distance between any selected two side-chain heavy atoms from the residues was less than 5Å (CZ of Arg, CG of His, NZ of Lys, CG of Asp, CD of Glu, OG of Ser, CB of Thr, CG of Asn, CD of Gln, SG of Cys, CA of Gly, CG of Pro, CB of Ala, CB of Val, CB of Ile, CG of Leu, CE of Met, CZ of Phe, OH of Tyr, CE2 of Trp). We selected only one heavy atom of the side chain to decrease the computational cost. The contacts in each frame of combined trajectories were calculated with Python using MDAnalysis^70^. A difference contact map was derived by subtracting down-state contacts from the up-state ones identified from the respective simulations.

All simulations were summarized in Table S3. Molecular visualizations were rendered using PyMOL^53^.

### Functional electrophysiology

#### Molecular biology

In this study the coding sequences of human K_2P_2.1 TREK-1 (Genbank accession number: NM_172042), rat K_2P_2.1 TREK-1 (NM_172041.2), human K_2P_10.1 TREK-2 (NM_021161) and human K_2P_4.1 TRAAK (AF247042) were used. For K^+^ channel constructs expressed in *Xenopus laevis* oocytes the respective K^+^ channel subtype coding sequences were subcloned into the dual-purpose vector pFAW. The TREK-2 construct with the extended pCt (TREK-2*) was truncated from WT and includes amino acids 67 - 355. All mutant channels (point mutations) were obtained by site-directed mutagenesis with custom oligonucleotides. Vector DNA was linearized with NheI or MluI and cRNA synthesized *in vitro* using the SP6 or T7 AmpliCap Max High Yield Message Maker Kit (Cellscript, USA) and stored at −20 °C (for frequent use) and −80 °C (for long term storage).

#### Inside-out patch-clamp measurements

*Xenopus laevis* oocytes were surgically removed from anesthetized adult females, treated with type II collagenase (Sigma-Aldrich/Merck, Germany) and manually defolliculated. A solution containing the K^+^ channel specific cRNA at a desired concentration was injected into Dumont stage V - VI oocytes and subsequently incubated at 17 °C in a solution containing (mM): 54 NaCl, 30 KCl, 2.4 NaHCO_3_, 0.82 MgSO_4_ x 7 H_2_O, 0.41 CaCl_2_, 0.33 Ca(NO_3_)_2_ x 4 H_2_O and 7.5 TRIS (pH 7.4 adjusted with NaOH/HCl) for 1 - 7 days before use. Patch-clamp recordings in inside-out configuration under voltage-clamp conditions were performed at room temperature (22 - 24 °C). Patch pipettes were made from thick-walled borosilicate glass GB 200TF-8P (Science Products, Germany), had resistances of 0.2 - 0.5 MΩ (tip diameter of 10 - 25 µm) and filled with a pipette solution (in mM): 120 KCl, 10 HEPES and 3.6 CaCl_2_ (pH 7.4 adjusted with KOH/HCl). Intracellular bath solutions and compounds were applied to the cytoplasmic side of excised patches for the various K^+^ channels via a gravity flow multi-barrel pipette system. Intracellular solution had the following composition (in mM): 120 KCl, 10 HEPES, 2 EGTA and 1 Pyrophosphate (pH adjusted with KOH/HCl). Currents were recorded with an EPC10 amplifier (HEKA electronics, Germany) and sampled at 10 kHz or higher and filtered with 3 kHz (−3 dB) or higher as appropriate for sampling rate.

#### Animals

The investigation conforms to the guide for the Care and Use of laboratory Animals (NIH Publication 85-23). For this study, twenty-five female *Xenopus laevis* animals were used to isolate oocytes. Experiments using Xenopus toads were approved by the local ethics commission.

#### Drugs

Tetra-pentyl-ammonium chloride (TPenA) was purchased from Merck (Sigma-Aldrich), prepared as 100 mM stocks in intracellular recording solution, stored at −20 °C and diluted to a final concentration in intracellular recording solution prior measurements. Decyl-MethaneThioSulfonate (Decyl-MTS), (2-(trimethylammonium)ethyl)MethaneThioSulfonate bromide (MTS-ET^+^) and Sodium (2-sulfonatoethyl)MethaneThioSulfonate (MTS-ES^-^) were purchased from Toronto Research Chemicals. MTS-ET^+^ was directly dissolved to the desired concentration of 1 mM in the intracellular recording solution prior to each experiment. MTS-ET^+^ was used immediately after dilution for maximally 5 min. Decyl-MTS and MTS-ET^-^ were prepared as 10 - 100 mM stocks in DMSO and diluted to the desired concentration prior the measurements. Norfluoxetine hydrochloride (NFx) was purchased from Cayman chemicals and prepared as 10 mM stocks in methanol.

### Data acquisition and statistics

Data analysis and statistics for functional electrophysiology were done using Fitmaster (HEKA electronics, version: v2×73.5, Germany), Microsoft Excel 2021 (Microsoft Corporation, USA) and Igor Pro 9 software (WaveMetrics Inc., USA).

Recorded currents were analyzed at voltage defined in the respective figure legend from stable membrane patches. Patch integrity was tested with a K_2P_ channel-specific blocker (1 mM TPenA).

The fold current increase (fold activation) of a ligand (decyl-MTS) was calculated from the following equation:

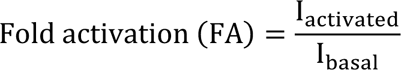

with I_activated_ represents the stable current level in the presence of 100 µM decyl-MTS and I_basal_ the measured current before application of the MTS probe.

The half-maximal concentration-inhibition relationship of a ligand (NFx) was obtained using a Hill-fit for dose-response curves as depicted below:

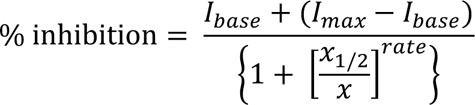

with base and max are the currents in the absence and presence of a respective ligand, x is the concentration of the ligand, x_1/2_ is the ligand concentration at which the inhibitory effect is half-maximal, rate is the Hill coefficient.

Data from individual measurements were normalized and fitted independently to facilitate averaging. Throughout the manuscript all values are presented as mean ± s.e.m. with n indicating the number of individual executed experiments. Error bars in all figures represent s.e.m. values with numbers (n) above indicating the definite number of executed experiments. A Shapiro-Wilk test or Kolmogorow-Smirnow test was used to determine whether measurements were normally distributed.

Image processing and figure design was done using Affinity Designer, Igor Pro 9 (64 bit) (WaveMetrics, Inc., USA) and Canvas X Draw (Version 20 Build 544) (ACD Systems, Canada).

## Data availability

Data supporting the findings of this manuscript are available from the corresponding authors upon request.

## Acknowledgements

This work was funded by the Deutsche Forschungsgemeinschaft (DFG) RU2518 DynIon (P1 to M.S. and T.B., P3 to H.S.) and the Leibniz-Forschungsinstitut für Molekulare Pharmakologie (FMP) (to H.S.). The MD simulations were performed with resources provided by the North-German Supercomputing Alliance (HLRN). We would like to acknowledge the help and support of Dr. Tillmann Utesch for helpful discussions.

