## Supplementary Tables S1-S3 for "From head to tail - Atomistic mechanism of long-range coupling from the cytosolic sensor domain to the selectivity filter in TREK K_2P_ channels"

### Table S1. Summary of apo TREK-2 computational electrophysiology simulations

| System | State | Model | PDB ID | $\Delta q$ ($e^-$) | Time (ns) | [C] (mM) | Vm (mV) | Ion permeations | $\gamma$ (pS) |
|---|---|---|---|---|---|---|---|---|---|
| TREK-2 | Up | Apo – Extended Length | 4BW5 | 2 | 1000 | 602.2 | 206 $\pm$ 10 | 5, 6, 3, 5, 6 | 4 $\pm$ 0.87 |
| TREK-2 | Up | Apo – Extended Length | 4BW5 | 2 | 1000 | 602.2 | -202 $\pm$ 7 | 0, 1, 2, 2, 2 | 1.12 $\pm$ 0.6 |
| TREK-2 | Down | Apo – Extended Length | 4XDJ | 2 | 1000 | 598.4 | 192 $\pm$ 12 | 0, 0, 0, 0, 0 | 0 |
| TREK-2 | Down | Apo – Extended Length | 4XDJ | 2 | 1000 | 598.4 | -189 $\pm$ 16 | 0, 0, 0, 0, 0 | 0 |

Each system contains 5 replicas. Temperature 300 K. $K^+$ and $Cl^-$ ions were used. AMBER99SB-ILDN force field and SPC/E water model.

**Table S2. Contacts from the interaction network analysis. The probability was calculated as the population difference between up-state and down-state contacts. If not specified as XXX$^{B}$, the residues come from chain A.**

| Intermolecular | | | Intramolecular | | | Intramolecular | | |
|---|---|---|---|---|---|---|---|---|
| Residue 1 | Residue 2 | Population difference (up – down) | Residue 1 | Residue 2 | Population difference (up – down) | Residue 1 | Residue 2 | Population difference (up – down) |
| S331$^{B}$ | F202 | 0.698 | A318 | F244 | 0.956 | G312 | L279 | 0.323 |
| K186$^{B}$ | F287 | 0.685 | D286 | Y273 | 0.899 | F252 | F215 | 0.317 |
| K114$^{B}$ | S150 | 0.648 | N149$^{B}$ | S145$^{B}$ | 0.889 | Y273 | F284 | 0.281 |
| A111$^{B}$ | S150 | 0.612 | A318$^{B}$ | F244$^{B}$ | 0.864 | F316 | T280 | 0.259 |
| A111$^{B}$ | S151 | 0.595 | Y315$^{B}$ | I245$^{B}$ | 0.855 | P303 | F163 | - 0.372 |
| A114$^{B}$ | D286 | 0.556 | S154 | N152 | 0.758 | R237 | K224 | - 0.409 |
| I110$^{B}$ | S150 | 0.504 | A247$^{B}$ | F215$^{B}$ | 0.751 | S181 | A142 | - 0.409 |
| F316$^{B}$ | P198 | 0.430 | D325 | R237 | 0.693 | K302$^{B}$ | H156$^{B}$ | - 0.415 |
| S155$^{B}$ | P101 | 0.404 | W326 | T241 | 0.684 | A289$^{B}$ | Y175$^{B}$ | - 0.443 |
| F316$^{B}$ | I197 | 0.399 | D325$^{B}$ | R237$^{B}$ | 0.672 | C189$^{B}$ | F165$^{B}$ | - 0.448 |
| I323$^{B}$ | P198 | 0.339 | Y315$^{B}$ | C249$^{B}$ | 0.652 | V288$^{B}$ | Y175$^{B}$ | - 0.452 |
| P180$^{B}$ | F284 | 0.297 | I170 | G167 | 0.627 | A293$^{B}$ | S145$^{B}$ | - 0.454 |
| L320$^{B}$ | P198 | 0.249 | F252$^{B}$ | F215$^{B}$ | 0.627 | F244 | S218 | - 0.490 |
| P198$^{B}$ | L320 | 0.142 | F316$^{B}$ | T280$^{B}$ | 0.608 | A318 | G212 | - 0.517 |
| P180$^{B}$ | E99 | - 0.247 | I178$^{B}$ | W157$^{B}$ | 0.567 | H156$^{B}$ | G143$^{B}$ | - 0.542 |
| C189$^{B}$ | Y273 | - 0.321 | G312$^{B}$ | L279$^{B}$ | 0.562 | R227 | E223 | - 0.557 |
| A111$^{B}$ | N152 | - 0.350 | F244$^{B}$ | G212$^{B}$ | 0.561 | F244$^{B}$ | S218$^{B}$ | - 0.567 |
| E270$^{B}$ | K186 | - 0.354 | S240$^{B}$ | E323$^{B}$ | 0.553 | G312 | F252 | - 0.585 |
| D118$^{B}$ | Q106 | - 0.392 | Q230 | R227 | 0.531 | I236$^{B}$ | F226$^{B}$ | - 0.601 |
| R328$^{B}$ | D325 | - 0.415 | A247 | F215 | 0.511 | I236 | V231 | - 0.635 |
| G91$^{B}$ | Y192 | - 0.447 | T278$^{B}$ | Y175$^{B}$ | 0.477 | G285$^{B}$ | G176$^{B}$ | - 0.651 |
| S155$^{B}$ | K107 | - 0.466 | F252 | L211 | 0.474 | S151 | S145 | - 0.685 |
| N141$^{B}$ | D286 | - 0.590 | I239$^{B}$ | F226$^{B}$ | 0.464 | R237$^{B}$ | E323$^{B}$ | - 0.713 |
| | | | F252$^{B}$ | L211$^{B}$ | 0.456 | I239 | F226 | - 0.739 |
| | | | F244 | G212 | 0.452 | W326$^{B}$ | R237$^{B}$ | - 0.794 |
| | | | K302 | H156 | 0.442 | Y297 | N292 | - 0.820 |
| | | | Y192 | I170 | 0.419 | G196 | I170 | - 0.836 |
| | | | A314 | G208 | 0.414 | Y297$^{B}$ | E264$^{B}$ | - 0.836 |
| | | | F284$^{B}$ | Y273$^{B}$ | 0.408 | Y297 | E264 | - 0.889 |
| | | | H156 | G143 | 0.380 | Y315$^{B}$ | A247$^{B}$ | - 0.893 |
| | | | T281 | T172 | 0.369 | A318$^{B}$ | L211$^{B}$ | - 0.920 |
| | | | G291 | E264 | 0.360 | S321$^{B}$ | G208$^{B}$ | - 0.936 |

**Table S3. Summary of all computational electrophysiology simulations**

| System | State | Model | PDB ID | Time per replica (ns) | Replica |
|---|---|---|---|---|---|
| (1) TREK-2 | Up | Apo – Extended Length | 4BW5 | 1000 | 5 |
| (2) TREK-2 | Down | Apo – Extended Length | 4XDJ | 1000 | 5 |
| (3) TREK-2 | Up | Phosphorylation – Extended Length | 4BW5 | 1000 | 5 |
| (4) TREK-2 | Down | Decyl-MTS – Extended Length | 4XDJ | 1000 | 5 |
| (5) TREK-2 | Down | MTS-$ET^+$ – Extended Length | 4XDJ | 1000 | 5 |

Temperature 300K. $K^+$ and $Cl^-$ ions were used. AMBER99SB-ILDN force field and SPC/E water model.
