## Supplementary Figures S1-S11 for "From head to tail - Atomistic mechanism of long-range coupling from the cytosolic sensor domain to the selectivity filter in TREK K_2P_ channels"

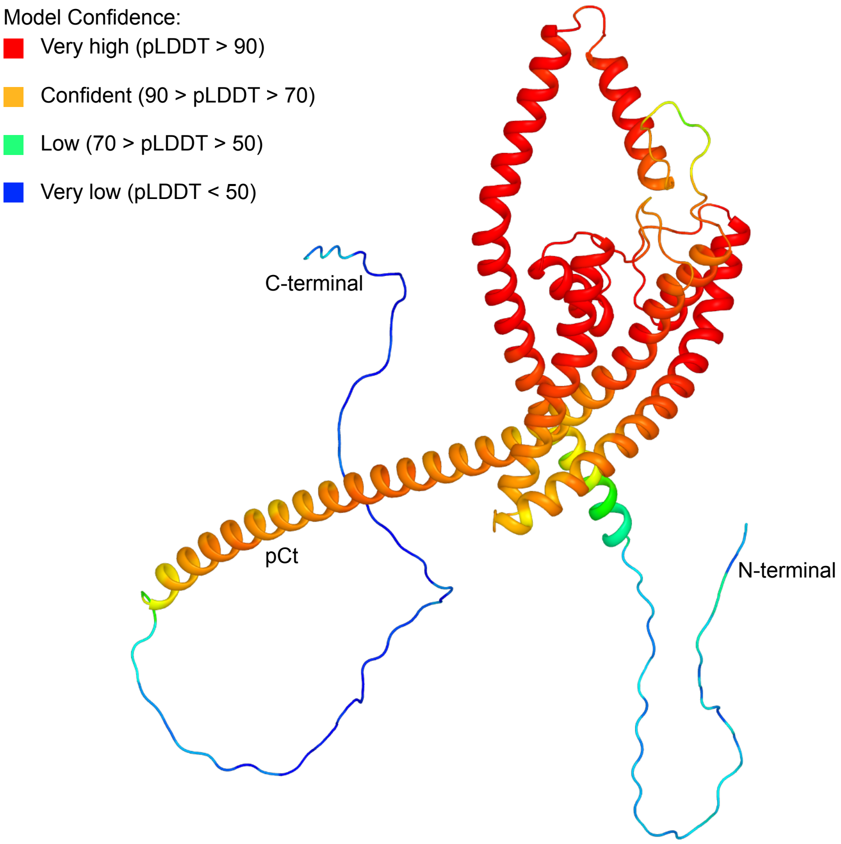


**Fig. S1. Predicted structure of KCNK2 (TREK-1) channel using AlphaFold2[1].**

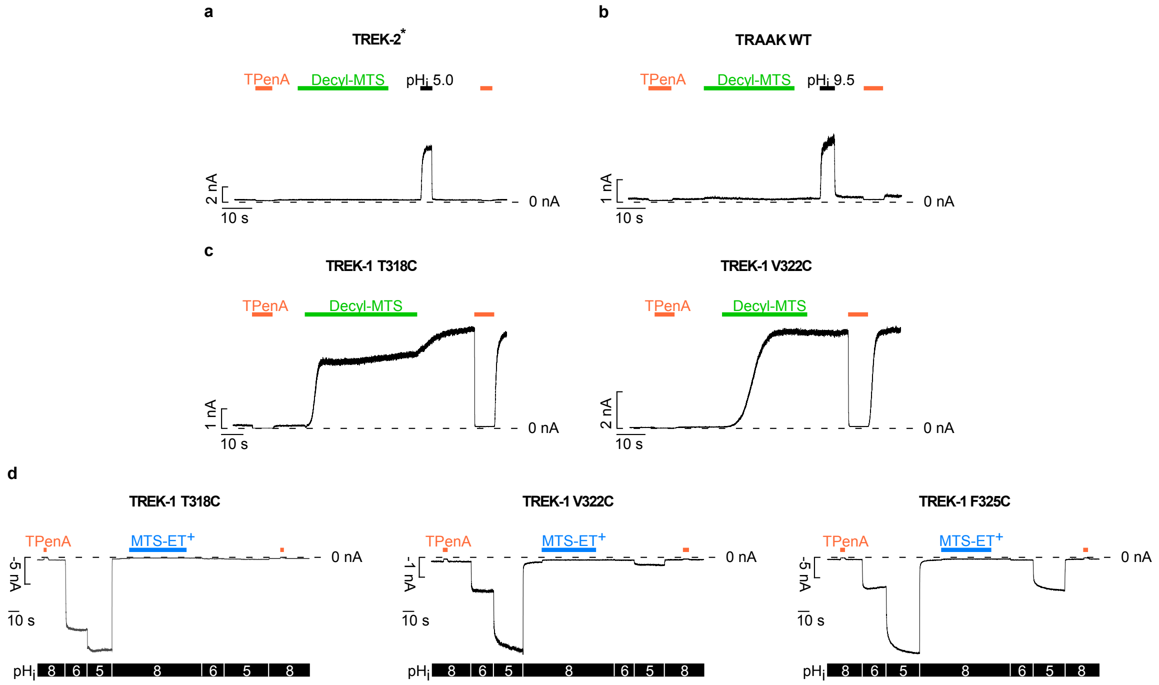


**Fig. S2. Validation of membrane tethering assay with cysteine modifying probes. (a,b)** Example recordings of TREK-2* (a) and WT TRAAK channels (b) in a symmetrical $K^+$ gradient at +40 mV and pH 7.4 showing robust activation with intracellular pH 5.0 or 9.5, respectively but no effect on the application of 100 µM decyl-MTS. **(c)** Same recordings as in (a,b) for T318C and V322C mutant TREK-1 channels showing strong and irreversible activation with 100 µM decyl-MTS. **(d)** Example recordings in a symmetrical $K^+$ gradient at -80 mV for T318C, V322C and F325C mutant TREK-1 channels showing robust activation upon intracellular acidification and channel inhibition with 1 mM MTS-$ET^+$.

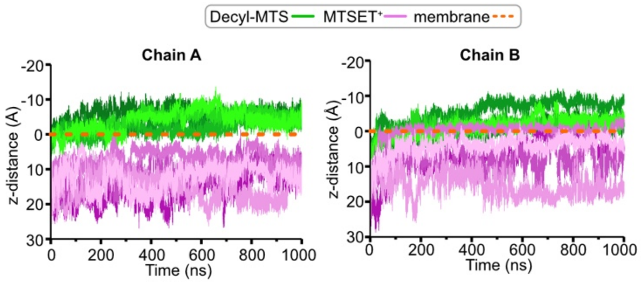


**Fig. S3. Distance between MTS ligand and the membrane derived from the simulations of MTS-modified TREK-2* simulations under -200 mV transmembrane voltages.** The distance (Dz component of the vector) between a chosen atom of the ligand (C12 for decyl-MTS, N2 for MTS-ET$^{+}$) and the membrane plotted against simulation time. The membrane was defined as a plane by selecting the center of geometry of phosphorus atoms located in the cytosolic part of the POPC lipid bilayer.

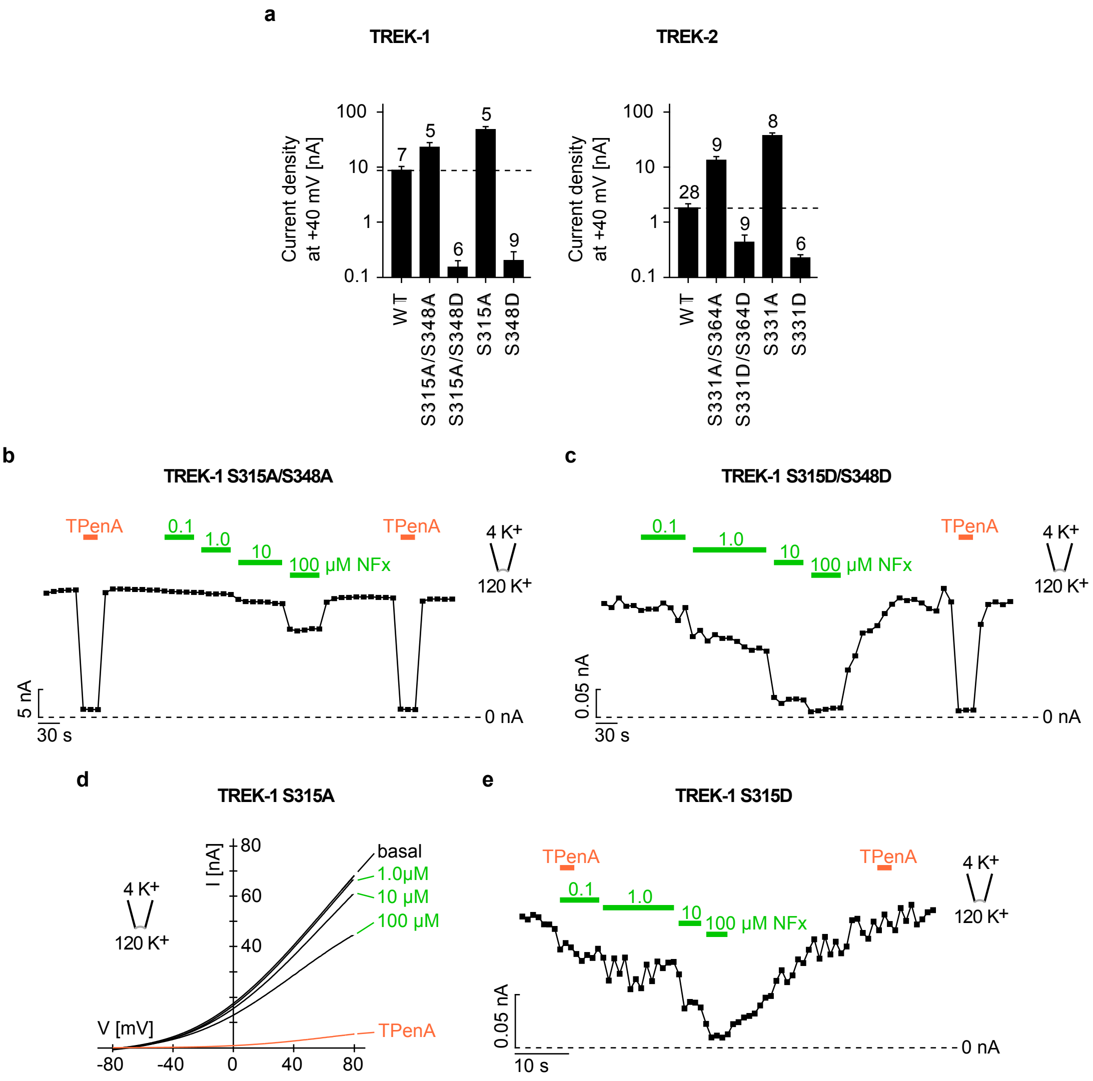


**Fig. S4. NFx sensitivity of TREK-1 (de-)phosphorylation mimicking mutant channels. (a)** Current densities of WT and mutant TREK-1/-2 channels illustrating the gain-of-function or loss-of-function effect of the (de-)phosphorylation mimicking mutations. **(b,c)** Analysis at +40 mV from measurements as in (d) for S315A/S348A (b) or S315D/S348D double mutant TREK-1 channels (c) showing an altered apparent affinity of NFx in contrast to WT. **(d)** Example recording of dephosphorylation at PKC site mimicking S315A mutant TREK-1 channels in an asymmetrical $K^+$ showing a decreased dose-dependent inhibition with NFx with indicated concentrations (green traces) and an almost full block with 1 mM TPenA (orange trace). (**e**) Analysis at +40 mV from measurements as in (d) for S315D mutant TREK-1 channels showing an altered apparent affinity of NFx in contrast to WT.

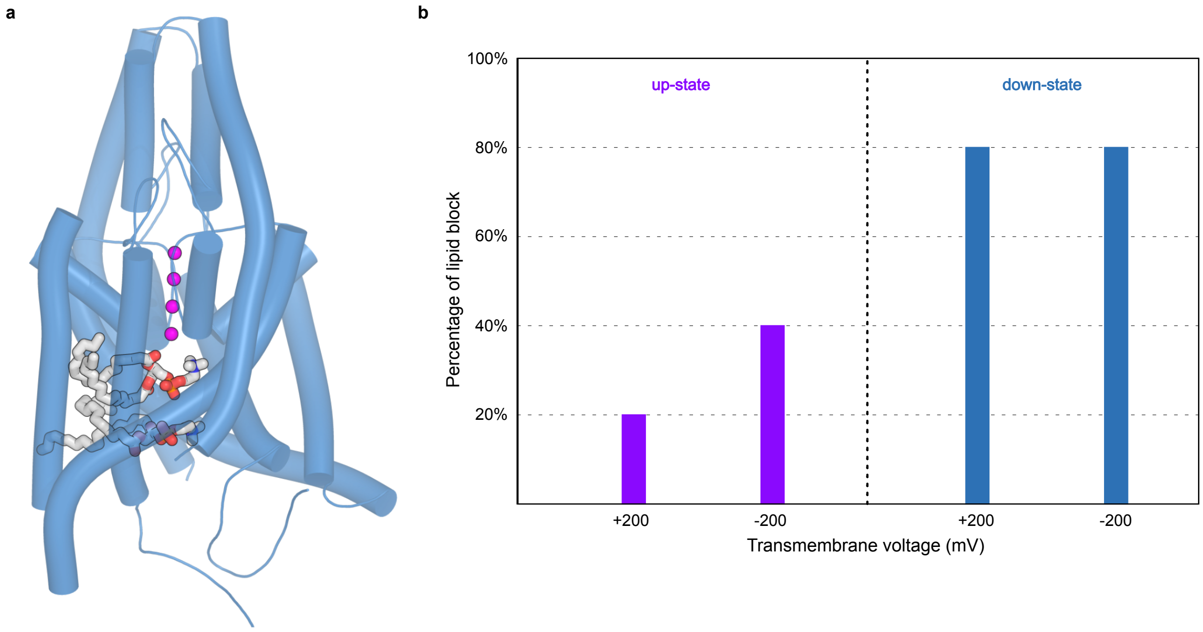


**Fig. S5. Lipid blockage of the inner pore**. **(a)** Selected end-snapshot from down-state simulations showing SF with $K^+$ ions (magenta) and the lipids (gray) blocking the fenestration side and the pore region. **(b)** Percentage of the simulations showing a lipid occupation in the inner pore of the TREK-2* channel during simulations. End-snapshots of each replica were used to analyze the number of blockage events.

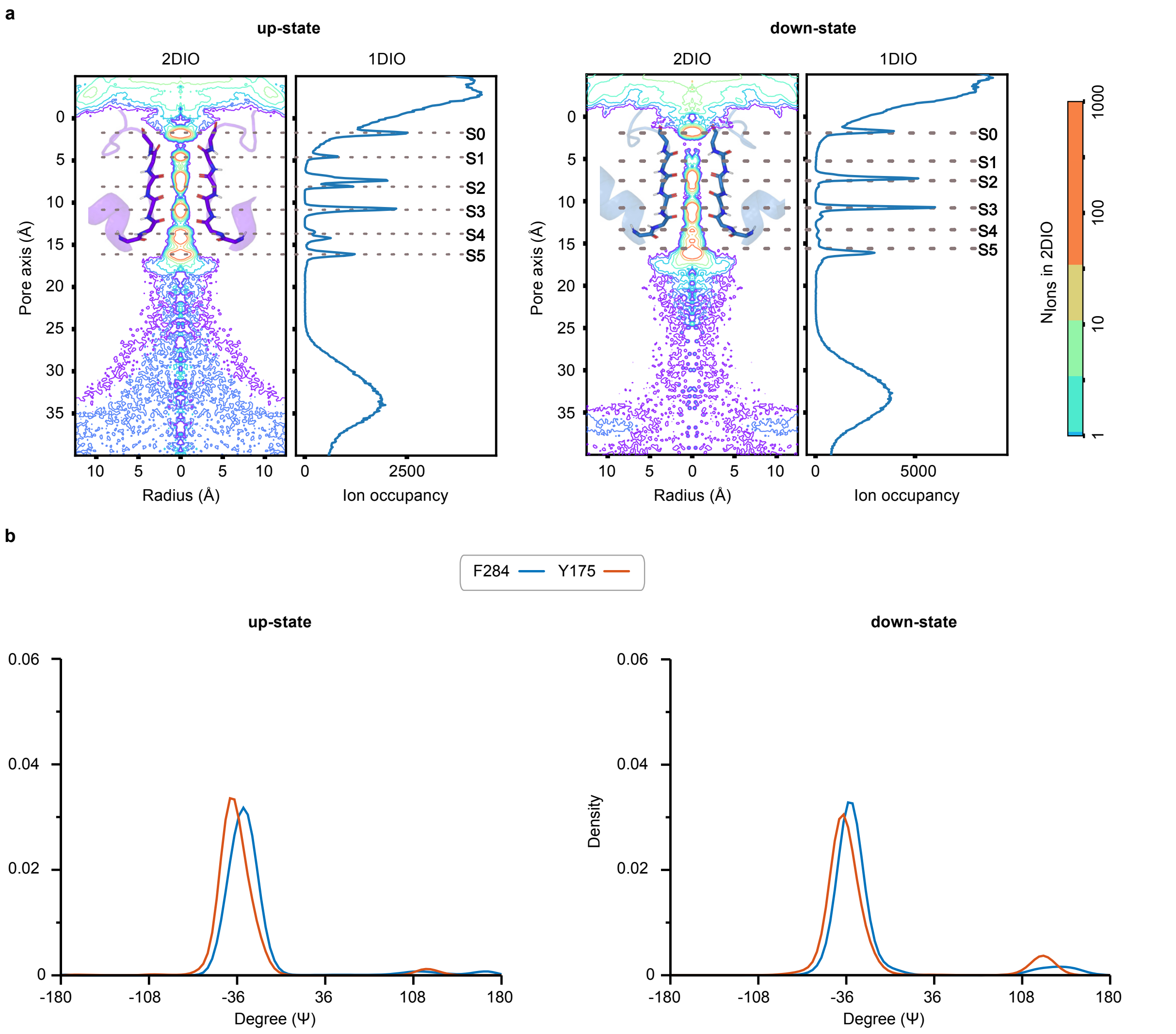


**Fig. S6. One- and two-dimensional ion occupancies. (a)** One- and two-dimensional ion occupancy within the SF for combined up-state (left) and non-conductive down-state (right) TREK-2* simulations at -200 mV. The radial area of the pore was defined and ions passing along the pore axis ($D_z$) were calculated from the simulations. Based on the volume change along the radius, the occupancy of ions was normalized per 0.001 Å$^3$ per 1 μs. The center of mass of the SF backbone atoms was located on the 7 Å point along the pore axis. Ion binding sites along the pore axis were indicated with dashed lines. **(b)** Psi angle (ψ) distributions of F284 and Y175 residues from the combined up-state (left) and down-state (right) simulations at -200 mV transmembrane potential.

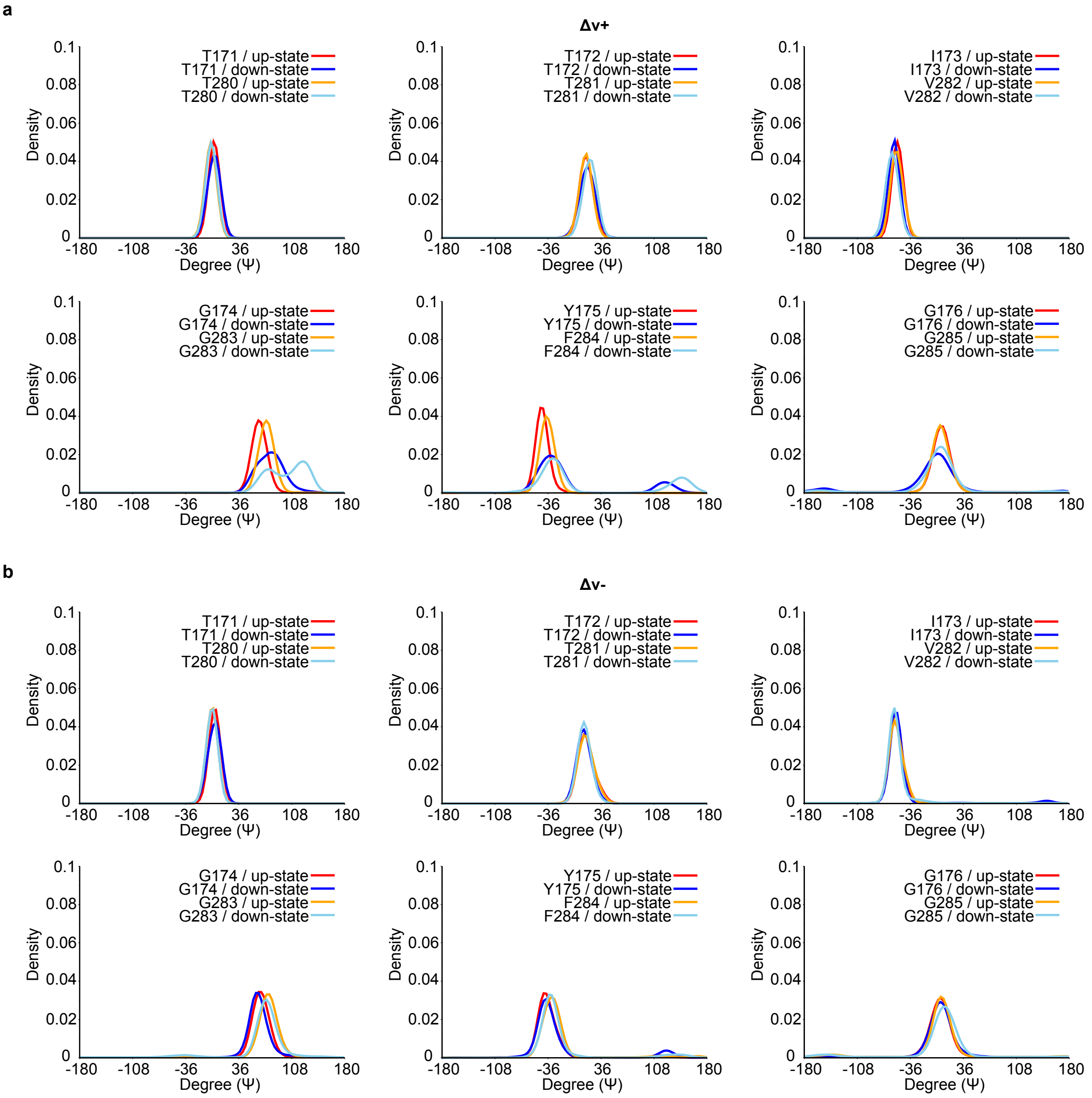


**Fig. S7. Conformational analysis of the SF residues.** Psi angle (ψ) distributions of SF residues from the combined up-state and down-state TREK-2* simulations at (a) +200 mV and (b) -200 mV transmembrane potential.

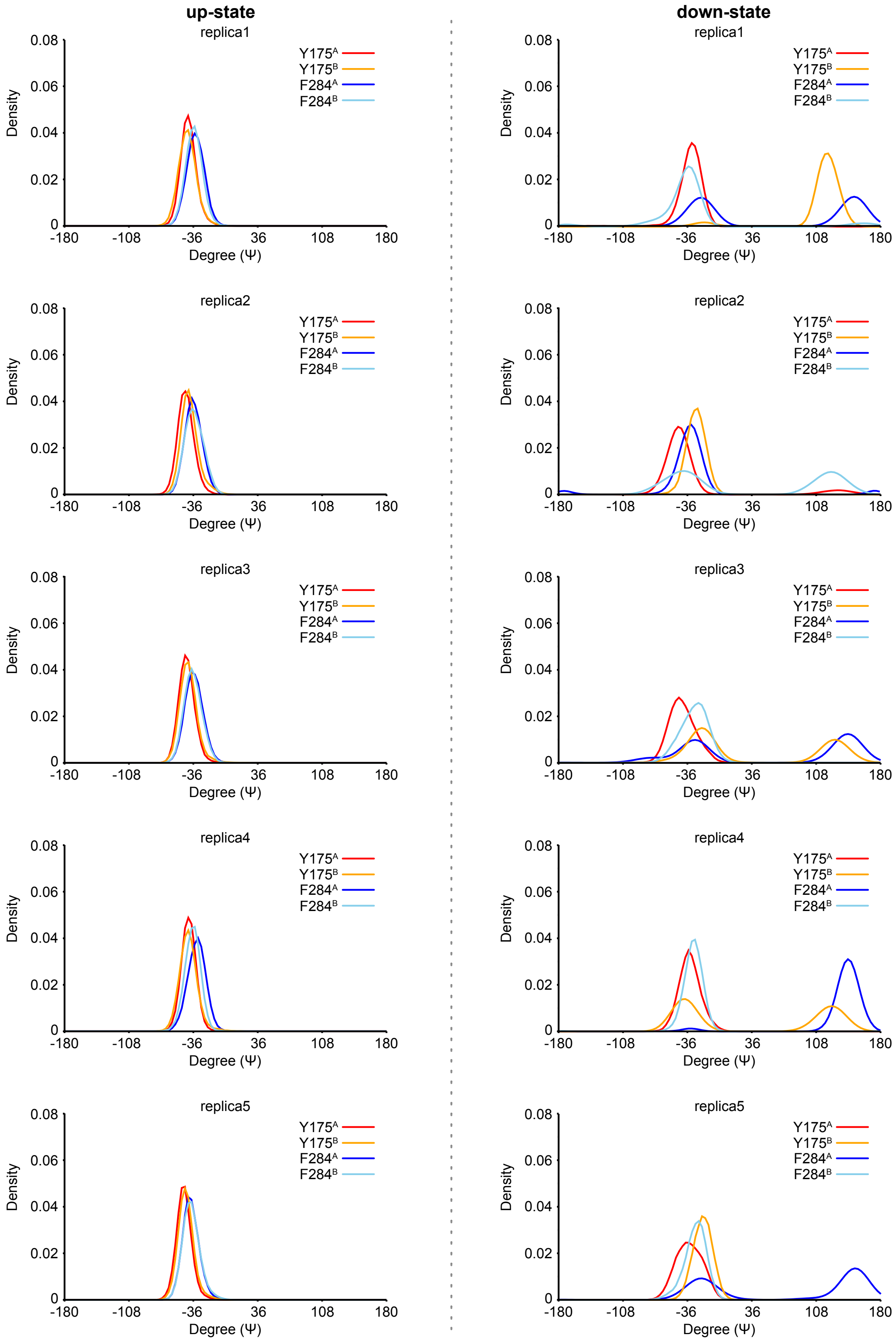


**Fig. S8. Conformational analysis of residues near S1 ion binding site.** Psi angles ($\psi$) of F284 and Y175 in chain A and chain B over the time from the up-state TREK-2* simulations at +200 mV transmembrane potential (left) and non-conductive down-state TREK-2* simulations at +200 mV transmembrane potential (right). Individual simulation runs were carried out with AMBER99sb[2] for 1 µs.

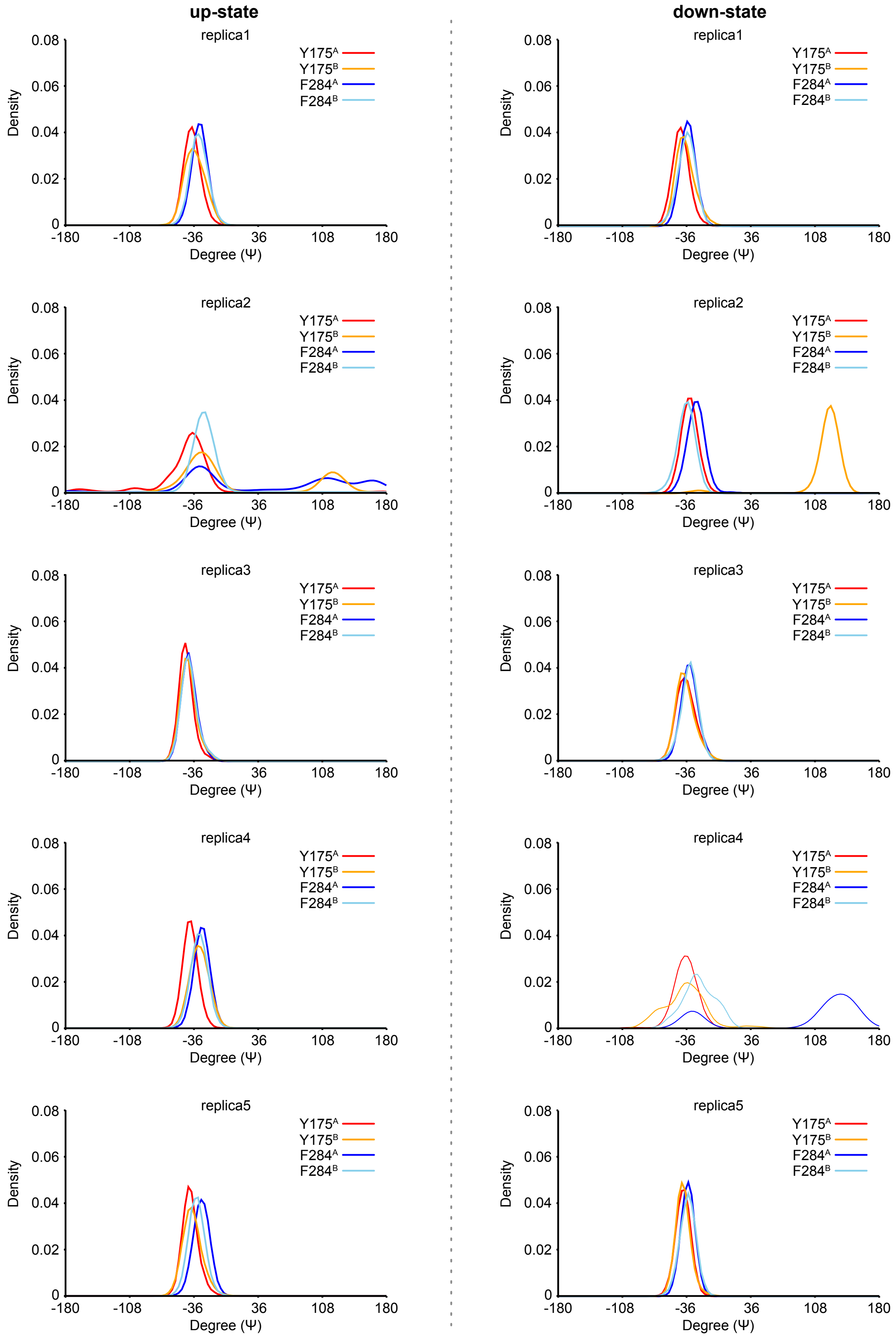


**Fig. S9. Conformational analysis of residues near S1 ion binding site.** Psi angles ($\psi$) of F284 and Y175 in chain A and chain B over the time from the up-state TREK-2* simulations at -200 mV transmembrane potential (left) and non-conductive down-state TREK-2* simulations at -200 mV transmembrane potential (right). Individual simulation runs were carried out with AMBER99sb[2] for 1 μs.

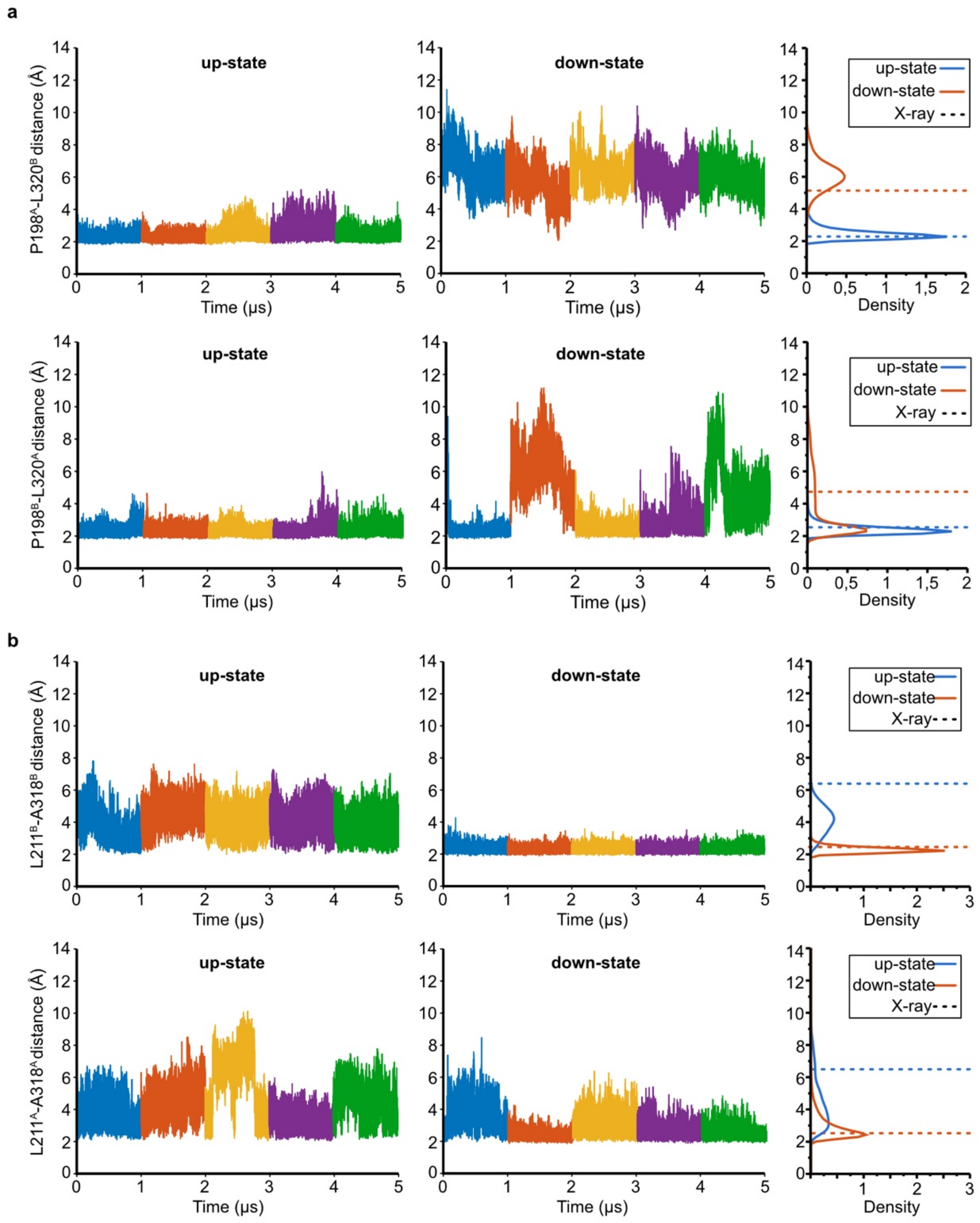


**Fig. S10. (a)** Time course of the inter-chain distance between P198 and L320 and **(b)** the intra-chain distance between L211 and A318 for both up- and down-state simulations of TREK-2*. Histograms of distance distribution are included on the right site, along with the distances derived from X-ray structures shown as dashed lines.

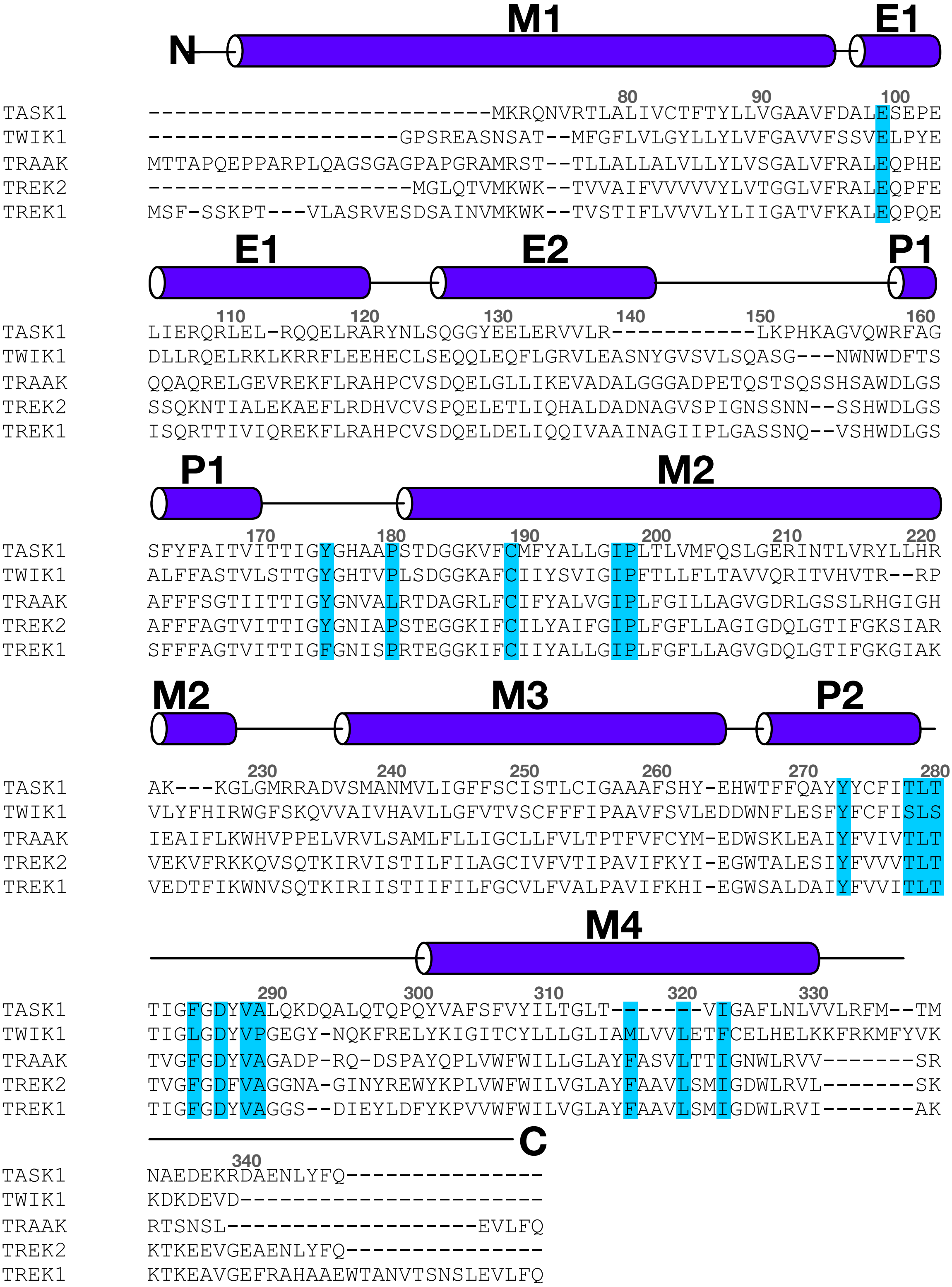


**Fig. S11. Sequence alignment of several K2P channels.** Sequence alignment of TASK-1 (PDB ID: 6RV2)[3], TWIK-1 (PDB ID: 3UKM)[4], TRAAK (PDB ID: 4WFE)[5], TREK-2 (PDB ID: 4BW5)[6] and TREK-1 (PDB ID: 6CQ6)[7] was generated by CLUSTALW2[8]. Important amino acid residues identified from the interaction network analysis are highlighted in blue.
